# Isolation of rat and human hepatic cholangiocytes using a peptide derived from a conserved domain of enterobacteria BamA/TamA-like proteins

**DOI:** 10.1101/2025.11.03.686257

**Authors:** Perla Metlej, Catherine Ribault, Sandrine David-Le Gall, Hanadi Nahas, Patricia Leroyer, Kevin Maunand, Nina Soler, Hugo Coppens-Exandier, Manuel Vlach, Dani Osman, Latifa Bousarghin, Anne Corlu, Pascal Loyer

## Abstract

Using the phage display technology, we identified a novel peptide, P11^Chol^, which preferentially binds to both human and rat cholangiocytes. Peptide P11^Chol^ alignment with protein databases evidenced strong similarities with a highly conserved peptide motif from BamA/TamA-like outer membrane proteins expressed in *enterobacteriaceae* belonging to Pseudomonadota phylum including *Photorhabdus*, *Providencia*, *Acinetobacter*, *Salmonella enterica* and *Helicobacter pylori* species. In addition, we showed that *Providencia stuartii* bacteria were able to bind to cholangiocytes-like HepaRG cells *in vitro* and that P11^Chol^ modulated this interaction suggesting the possible involvement of BamA/TamA-like outer membrane proteins in cell adhesion and/or internalization of *Providencia stuartii* bacteria. Using fluorescent P11^Chol^ peptide, we next developed a flow cytometry procedure to detect and isolate rat and human liver epithelial cells from hepatic cell suspension obtained after collagenase dissociation of liver parenchyma. Three distinct P11^Chol^-positive rat liver epithelial cell lines (RLEC) were established, which produced functional cholangiocytes capable to form cyst-like structures *in vitro* and to maintain expression of specific functions in hepatocytes in coculture. The characterization of these three RLEC lines evidenced functional differences that support the concept of small and large cholangiocytes exhibiting different functional phenotypes within the intrahepatic bile tree.

## 1. Introduction

Antibody-based therapies represent the dominant “biologics” in medicine [1], however, many small proteins and peptides have been approved for clinical applications [2] and novel therapeutic peptides are currently under clinical trials, which should expand their use in a near future for innovative therapeutics [3, 4, 5]. The first peptide-based drugs were used for treatment of endocrine and metabolic disorders with naturally occurring or synthetic peptides among which insulin, glucagon-like peptide-1 agonists, growth hormone and somatostatin are emblematic of peptide-biologics in medical practice [3]. More recently, peptides have also been developed for many other medical applications including non-opioid analgesics [6], antiviral bioproducts [7, 8], antibiotics [9, 10, 11] as well as antihypertensive drugs and cardioprotective agents [5]. Furthermore, recent advances in plasma peptidomics opened the perspective to identify biomarkers for the diagnosis and stratification of diseases [12, 13].

Peptides have also garnered growing interest as cell targeting ligands since they exhibit specific binding to plasma membrane receptors. In this context, peptide-drug conjugates [14, 15, 16, 17], radiolabeled peptides for tumor imaging and therapy [18, 19, 20, 21] and anticancer peptides inducing selective cytotoxicity in cancer cells [22] have been developed. Similarly, nanoparticles (NPs) are functionalized with peptide ligands of membrane receptors expressed in tumor cells in order to enhance drug delivery within tumors [14, 23, 24, 25, 26] although the efficacy of this strategy remains controversial [27, 28]. Peptides have also been modified with fluorochromes in order to label specific cell types for various biotechnological applications including cell isolation and adhesion [29, 30, 31] and phenotyping [32, 33].

Novel bioactive and/or cell-targeting peptides are identified by different methodological approaches including purifications and proteomic/mass spectrometry analysis of naturally occurring peptides in biological fluids or cell extracts [13]. The phage display technology also allows the isolation of short peptides from collections of random sequences introduced into the genome of a phage surface protein [34]. Recombinant phage libraries incubated in presence of cell targets lead to the selection of candidate amino acid motifs with putative binding capacity to a given cell type, which can be further validated using synthetic peptides. This technology is also used to characterize peptide epitopes of antibody clones [35].

The phage display has been successfully used to identify peptides interacting with various cell types including human and murine hepatoma cells [36–46], LX2 hepatic stellate myofibroblasts [47] and the liver macrophages [48]. In contrast, only one study has described phage display screen using human intrahepatic cholangiocarcinoma-derived RBE cells [49] to identify peptide COP35 as a binding partner of the clathrin heavy chain associated to glucose-regulated protein of 78 kDa (GRP78/BiP) [50].

In this study, we used the human cholangiocyte-like HepaRG cells to perform a phage display screen in order to identify novel(s) peptide(s) targeting cholangiocytes. In contrast with transformed cholangiocyte-like cells such as RBE [49], the HepaRG cells do not originate from an intrahepatic cholangiocarcinoma but are bipotent hepatic progenitors isolated from an hepatocellular carcinoma (HCC) capable to differentiate into both biliary- and hepatocyte-like cells in appropriate culture conditions [51–54]. In addition, HepaRG cells have kept the major cell cycle check-points since they exhibit wild-type *P53*, *RB*, *WNT* and *β-catenin* genes, which contribute to their long-term stability when cell monolayers of differentiated cholangiocyte and hepatocyte-like cells are kept confluent over several weeks [51–54].

We took advantage of the cholangiocyte-like HepaRG cells of as cell baits to identify the novel peptide P11^Chol^, which showed strong similarities with a highly conserved peptide motif from BamA/TamA-like outer membrane proteins expressed in *enterobacteriaceae*. Using a fluorescent peptide P11^Chol^, we developed the detection by flow cytometry of rat and human intrahepatic liver epithelial cells from hepatic cell suspensions. The present study provides new means to isolate primary liver epithelial cells.

## 2. Results

### 2.1. Peptide P11^chol^ strongly binds to Cholangiocyte-like HepaRG cells and RLEC

The phage display screen was performed using the hepatocyte-like HepaRG and HepG2 cells to deplete nonspecific phages (**Supporting Information-SI 1**). Then, supernatant containing phages that did not bind to these cells were incubated with the Chol^DMSO^-HepaRG cells prior to the elution of the bound phages. Thirty-three clones encoding 16 different full-length peptides were identified (**SI 2B**). The affinity of these 16 clones of phages towards the Chol^DMSO^-HepaRG cells was further evaluated in a colorimetric ELISA-like assay. The clones 10, 11, 12, 18, 46, 50 and 51 that showed an optical density above the baseline were selected for the next experiments (**SI 2C**).

The peptides corresponding to the selected phage clones were synthesized (**SI 3A**) with a biotin residue covalently bound to the C-terminus for *in vitro* cell-binding assay using fluorescent Dylight^TM^488 streptavidin (**SI 3B**). We first evaluated the binding of the peptide-streptavidin complexes to Chol^DMSO^-HepaRG cells at 3 different concentrations by measuring the mean of fluorescence and the percentages of positive cells (**Fig 1**). As negative controls, untreated cells and cells incubated with biotin-Dylight^TM^488 streptavidin were used. While Chol^DMSO^-HepaRG cells incubated with the peptide 10-, 12-, 18-, 50- and 51-Dylight^TM^488 streptavidin complexes showed similar fluorescence levels to those measured with negative control cells, more than 90% of the cells were labelled with the peptide 11 (**Fig 1A**), and mean fluorescence increased in a dose-dependent manner (**Fig 1B**). The peptide 46 also slightly increased the fluorescence (**Fig 1A, C**) but in a much weaker level compared to peptide 11. Independent experiments confirmed that peptide 11 strongly bound to Chol^DMSO^-HepaRG cells while the peptide 46 weakly interacted with these cells (**Fig 1D**).

**Figure 1.**
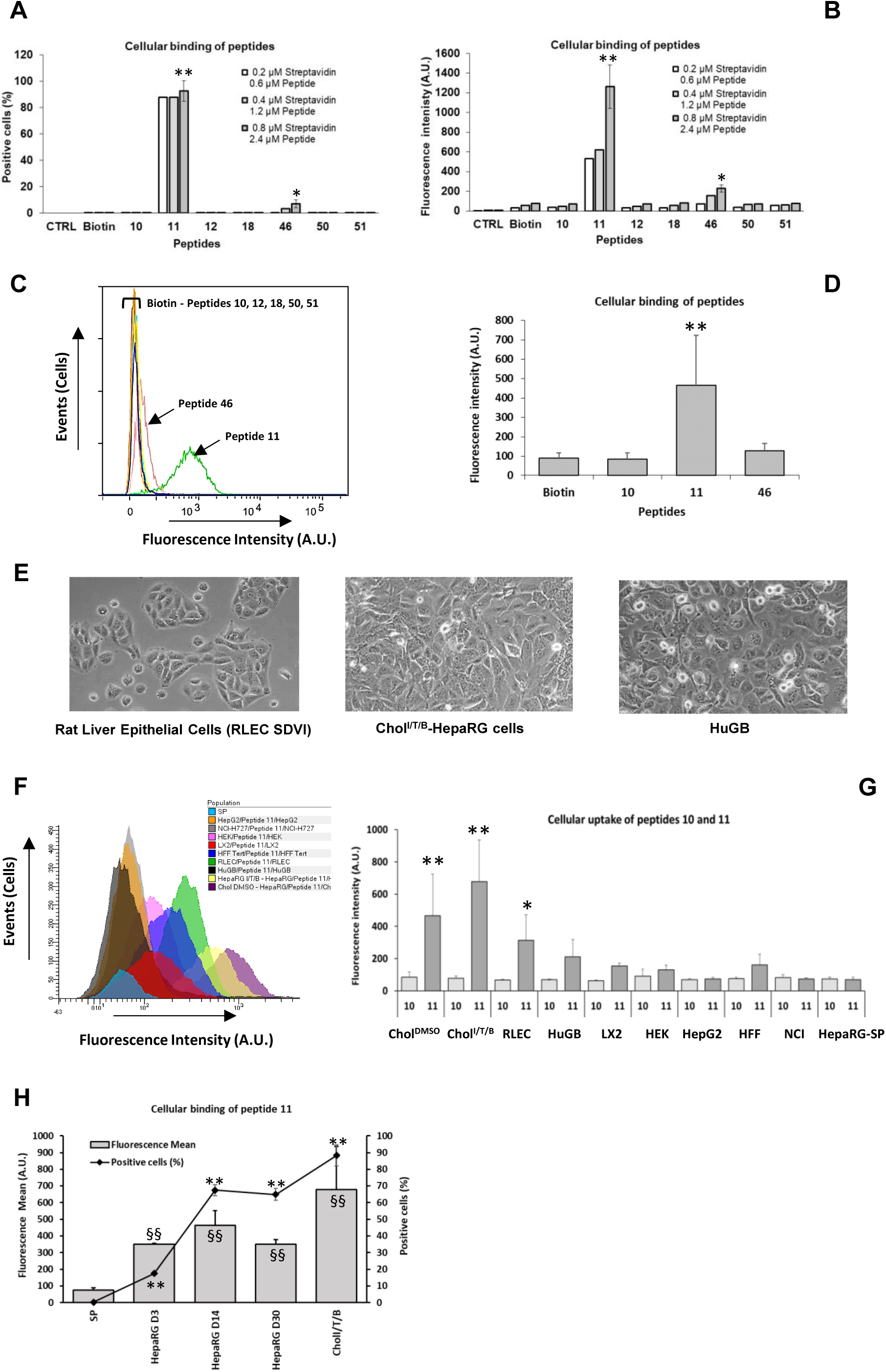
Peptide P11^Chol^ preferentially binds to Chol^I/T/B^-HepaRG cells, RLEC but not progenitor HepaRG and HepG2 hepatoma cells. Cell binding assay of synthetic peptide-fluorescent Dylight^TM^488 streptavidin complexes to Chol^DMSO^-HepaRG cells evaluated by flow cytometry and expressed as relative fluorescence intensity in arbitrary units (A) and percentage of positive cells (B) using different concentrations of peptides 10, P11^Chol^, 12, 18, 46, 50 and 51, and Dylight^TM^488 streptavidin compared to the negative control using Dylight^TM^488 streptavidin only (CTRL), or Dylight^TM^488 streptavidin bound to biotin only without peptide (Biotin). Concentrations in peptides and streptavidin were 0.2μM peptide/0.6μM streptavidin (2 independent experiments, n=2), 0.6μM peptide/1.8μM streptavidin (2 independent experiments, n=2), and 0.8μM peptide/2.4μM streptavidin (3 independent experiments, n=3). C) Superposition of the cytometry histograms showing the fluorescence shift obtained with P11^Chol^ in a typical experiment. D) Evaluation of the binding of the peptides 10, P11^Chol^ and 46 to the Chol^DMSO^-HepaRG cells in 4 independent experiments (n=4). Statistics: *p<0.05 and **p<0.01 between peptide-fluorescent Dylight^TM^488 streptavidin complexes and control peptide 10-Dylight^TM^488 streptavidin conjugates. E) Phase contrast photographs showing the cell morphology of Rat Liver Epithelial Cells (RLEC), Chol^DMSO^- and Chol^I/T/B^ HepaRG cells, and HuGB cells. F) Overlay of monoparametric histograms of flow cytometry obtained after incubating P11^Chol^ with RLEC, Chol^DMSO^- and Chol^I/T/B^ HepaRG cells, and HuGB cells, HepG2, LX2, HepaRG-SP or non-hepatic cells (NCI-H727, Human Foreskin Fibroblasts-HFF, HEK293T). The different cell lines were incubated with a 0.8 μM peptide : 2.4 μM streptavidin ratio for 24h and the means of fluorescence of the cell populations were measured by flow cytometry. Separated histograms are provided in Supporting Information 7. G) Relative fluorescence quantifications of the Chol^DMSO^- and Chol^I/T/B^ HepaRG cells, RLEC, HuGB, LX2, HepG2, Human Forskin Fibroblasts (HFF), LX2, HEK293T, NCI-H727 and progenitor “side-population” HepaRG-SP following binding of peptides 10- and P11^Chol^-Dylight^TM^488 streptavidin complexes. H) Biparametric charts showing mean fluorescence (mean in arbitrary units, A.U.) and percentage of positive cells (%) after incubation of P11^Chol^-Dylight^TM^488 streptavidin complexes with the different HepaRG cells: progenitor “side-population” cells (SP), HepaRG proliferating progenitor cells 3 days after plating (HepaRG D3), quiescent cells undergoing differentiation at day 14 (HepaRG D14) and at day 30 (HepaRG D30) days after plating, and cholangiocyte Chol^I/T/B^ HepaRG cells. Statistics: C) **p<0.01 and ^§§^p<0.01 for percentages of positive cells and means of fluorescence, respectively, between HepaRG D14, D30 and progenitor “side-population” cells (SP).

We next investigated whether peptide 11, further referred as P11^Chol^, also bound to other cholangiocyte cell models (**Fig 1E**), the rat liver epithelial cells (RLEC SDVI) [55], the Chol^I/T/B^-HepaRG cells [52] and the HuGB [56], as well as the LX2 hepatic stellate, HepG2, HEK 293, NCI-H727 cells and the HepaRG Side Population (HepaRG-SP cells) corresponding to early hepatic progenitor cells [57]. High percentages of P11^Chol^ positive cells (>90%, data not shown) were found for the RLEC, Chol^I/T/B^- and Chol^DMSO^-HepaRG cells, with much higher fluorescence levels compared to those found in undifferentiated HepaRG-SP, LX2, HepG2, HEK 293 and NCI-H727 cells (**Fig 1F-H, SI 4**). P11^Chol^ bound to a small fraction of the HuGB cells (∼5%); which were strongly positive (**Fig 1G, SI 4**).

We also compared the binding of P11^Chol^-Dylight^TM^488 streptavidin complexes to HepaRG cells at the different stages of the differentiation in HepaRG cells (**Fig 1H**). P11^Chol^ bound to less than 1% of HepaRG-SP cells and to nearly 18% of the bipotent progenitor HepaRG cells 3 days after plating with a 4-fold increase in fluorescence intensity between these 2 culture conditions. The percentage of positive cells increased throughout the differentiation process. The intensities of fluorescence were not significantly different at day 30 in both hepatocyte- and Chol^DMSO^-HepaRG cells. For Chol^I/T/B^-HepaRG cells, the means of fluorescence were significantly higher than those measured in the other culture conditions. We visualized the internalization of P11^Chol^-Dylight^TM^488 streptavidin complexes by confocal microscopy in Chol^DMSO^-HepaRG cells and RLEC (**SI 5A**) and demonstrated their internalization by endocytosis using endocytosis inhibitors chlorpromazine and genistein (**SI 5B**).

### 2.2. P11^Chol^ is similar to membrane proteins expressed in bacteria

P11^Chol^ [seq: seq: TFLNSVPTYSYWGGGS] is a 1735.87 g/L theoretical molar weight peptide with GRAVY score of -0.175 and an isoelectric point at 5.18 (**SI 6**). Its amino acid sequence showed high scores of identities with a well-conserved domains within BamA/TamA-like outer membrane proteins from *Photorhabdus asymbiotica*, *Providencia stuartii*, *Acenitobacter baumanni*, *Salmonella enterica*, the tannase and feruloyl esterase expressed in *Alternaria alternata*, a Pebp-like protein from *Helicobacter Pylori*, the Hepatitis C virus polyprotein and an ABC transporter permease from *Ruthenibacterium lactatiformans* (**Table 1**), which delineated a consensus peptide motif [seq: L-N-(S/N)-(V/I)-P-T-Y-Y-W-G-X-G]. A partial alignment with the human E-selectin lectin was also evidenced. We extended the alignment on both ends of the consensus peptide motif with some of these bacterial proteins (**Fig 2A**) and demonstrated that Bam/Tam A-like proteins were highly similar (> 90%) while the others proteins exhibited weaker similarities, excepted within the consensus peptide motif similar to P11^Chol^. We next looked for the position of the consensus peptide motif within crystal structures of TamA-like proteins available for *Escherichia coli* and *Salmonella enterica* (**Fig 2B-C**). The TamA/Omp85 protein in *Escherichia coli* comprises a 16-stranded transmembrane β-barrel in which the tyrosine Y328 aligned with the second Tyrosine (Y11) of P11^Chol^ (**Fig 2B**). In the β-barrel BamA-B-C-D-E assembly machinery complex of *Salmonella enterica*, the same Y11 of P11^Chol^ aligned to Y230 of the BamA subunit C located in a helix domain facing the intracellular end of the β-barrel (**Fig 2C**). Since no crystal structures were available for the Tam/BamA-like proteins in *Providencia stuartii* bacteria, we used iCn3D software that predicted a transmembrane β-barrel in which the Y135 aligning with the Y11 of P11^Chol^ located within an extracellular motif of the β-barrel (**Fig 2D**). Together, these data identified a conserved peptide motif in the transmembrane β-barrel of Tam/BamA-like outer membrane proteins highly similar to P11^Chol^.

**Figure 2.**
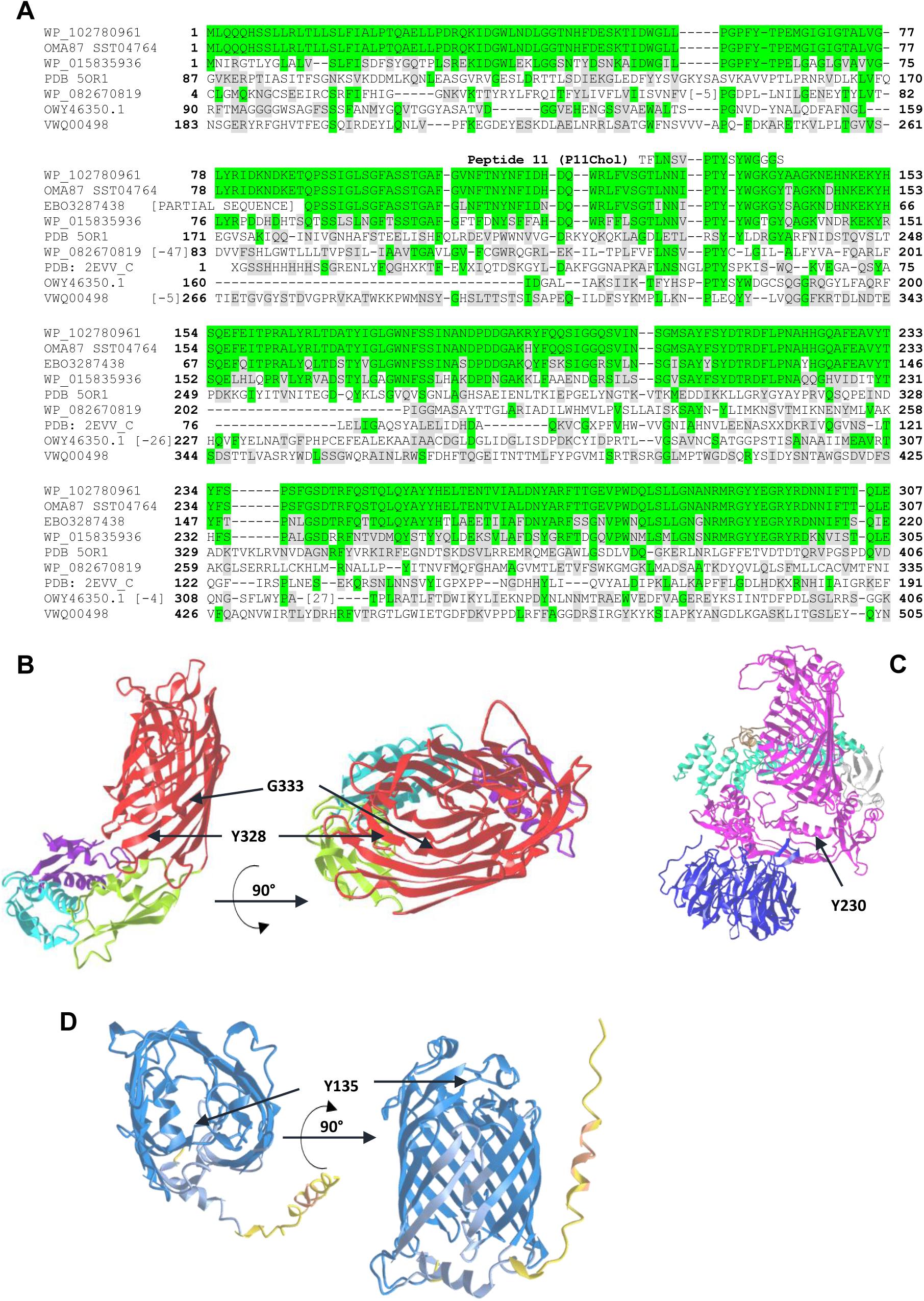
Amino acid sequence alignments of bacteria proteins containing a peptide motif similar to P11Chol and crystal structures of BamA/TamA-like proteins. A) Extended alignments of BamA/TamA-like proteins from *Providencia stuartii* (WP_102780961.1), *Acinetobacter baumannii*, (SST04764.1) *Salmonella enterica* serovar Enteritidis, (EBO3287438), *Photorhabdus asymbiotica*), (WP_015835936.1) tannase and feruloyl esterase (*Alternaria alternata*), (OWY46350.1) ABC transporter permease (*Ruthenibacterium lactatiformans*) (WP_082670819.1) and Pebp-like protein (*Helicobacter Pylori*) (PDB: 2EVV_C). Green squares show conserved amino acids between BamA/TamA-like proteins from *Providencia stuartii* and the other proteins. Grey squares represent conservative substitution. [X] symbol for the number of amino acids, indicated partial amino acid sequences not presented. Peptide 11 (P11Chol) is positioned to evidence the consensus peptide motif [L-N-(S/N)-(V/I)-P-T-Y-Y-W-G-X-G]. Numbers at each end of lanes indicate the positions of amino acids within the protein sequences. Crystal structures (iCn3D, NCBI) of B) TamA/Omp85 protein in *Escherichia coli* (PDB ID 4C00), in which Tyr328 aligning with the second Tyr of P11^Chol^ is located into the transmembrane β-barrel, C) Entire outer membrane β-barrel BamA-B-C-D-E assembly machinery complex from *Salmonella enterica serovar Typhimurium* (PDB ID 5ORI), with the position of Tyr230 aligning with the second Tyr of P11^Chol^. D) Predicted structure (iCn3D) of Bam/Tam A outer membrane protein of *Providencia stuartii* (WP_102780961.1) with transmembrane β-barrel containing Tyr135 aligning with the second Tyr of P11^Chol^.

**Table 1.**
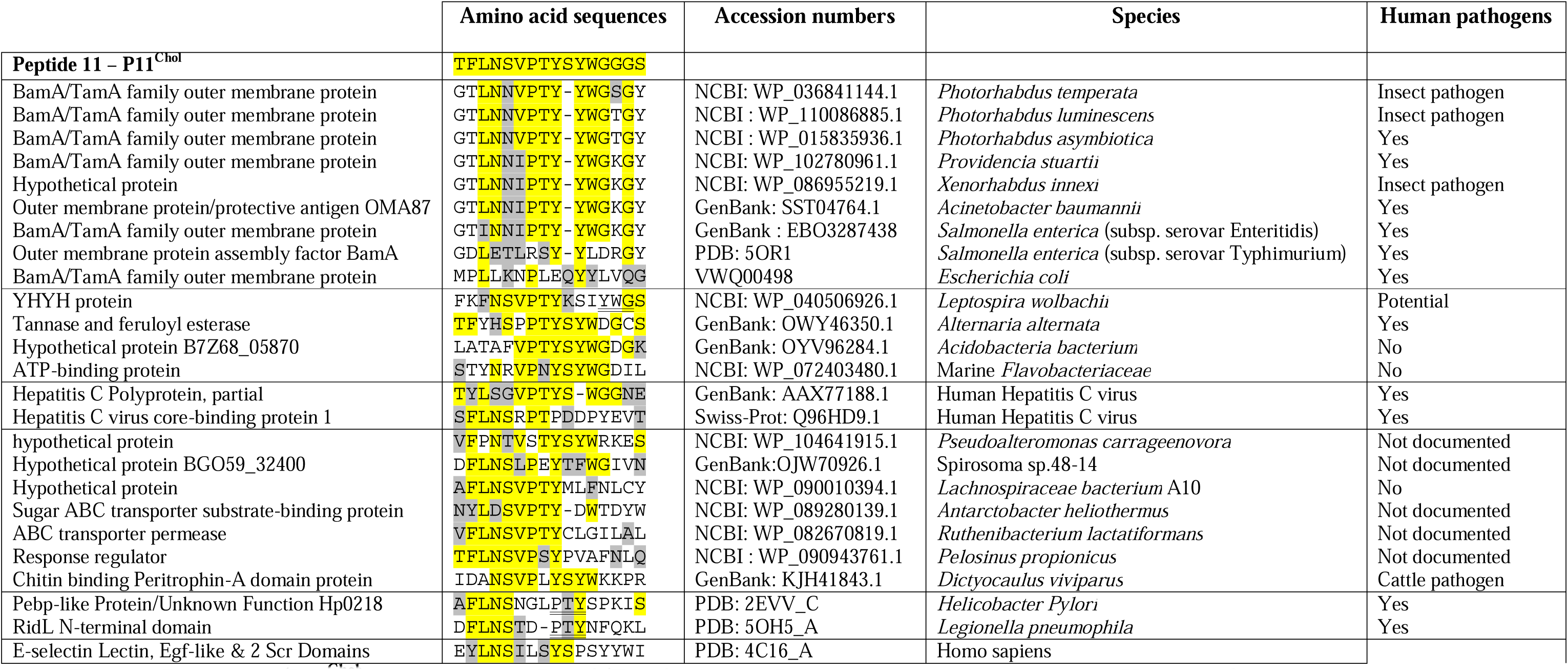
Peptide 11 (P11^Chol^) homology with BamA/TamA-like and Pebp-like protein from bacteria, Hepatitis C virus polyprotein and human E-selectin. Alignment of P11^Chol^ and peptides derived from membrane proteins of various bacteria species, hepatitis C virus (HCV) and human E-selectin. Alignments were performed using Cobalt at NCBI (NIH, USA) and refined manually. The amino acids highlighted in yellow are conserved, the related amino acids (substitution) are highlighted in gray. Accession numbers in protein data bases are provided. Scores of identities with BamA/TamA-like outer membrane proteins were >80% with 8 to 10 identical amino acids out of 16 and conservative substitutions.

### 2.3. Microbiota in healthy human livers and intrahepatic cholangiocarcinomas

We next analyzed the microbiota using 16S rRNA sequencing of the in healthy human livers, paired peritumoral livers and intrahepatic cholangiocarcinomas (CCK) to determine if bacteria expressing Tam/BamA-like proteins containing the consensus peptide motif similar to P11^Chol^ could be present in human livers (**Fig 3A**). Four main phyla were evidenced: Actinomycetota, Bacteroidota, Bacillota, Pseudomonadota and undetermined bacteria. Interestingly, we showed that the alpha-diversities were significantly higher in peritumoral parenchyma and CCK compared to healthy livers (**Fig 3B**) and we found that Actinomycetota and Bacilota phyla were significantly more abundant in CCK compared healthy and peritumoral livers while Pseudomonadota phylum abundance was reduced (**Fig 3C**). Several bacteria such as *Acinetobacter baumannii*, *Providencia stuartii*, *Helicobacter Pylori* and *Photorhabdus asymbiotica*, which express Tam/BamA-like proteins belong to the abundant phyla identified in the microbiota analysis, particularly the Pseudomonadota phylum. We also demonstrated by RT-qPCR the presence of 16S RNA from *Providencia*, *Photorhabdus* and *Salmonella enterica* species in total RNAs extracted from healthy and peritumoral livers and CCK (**Fig 3D**) demonstrating that both human normal livers and CCK contained bacteria belonging to these different phyla. In addition, these 16S RNA levels were significantly higher in peritumoral livers compared to the two other groups.

**Figure 3.**
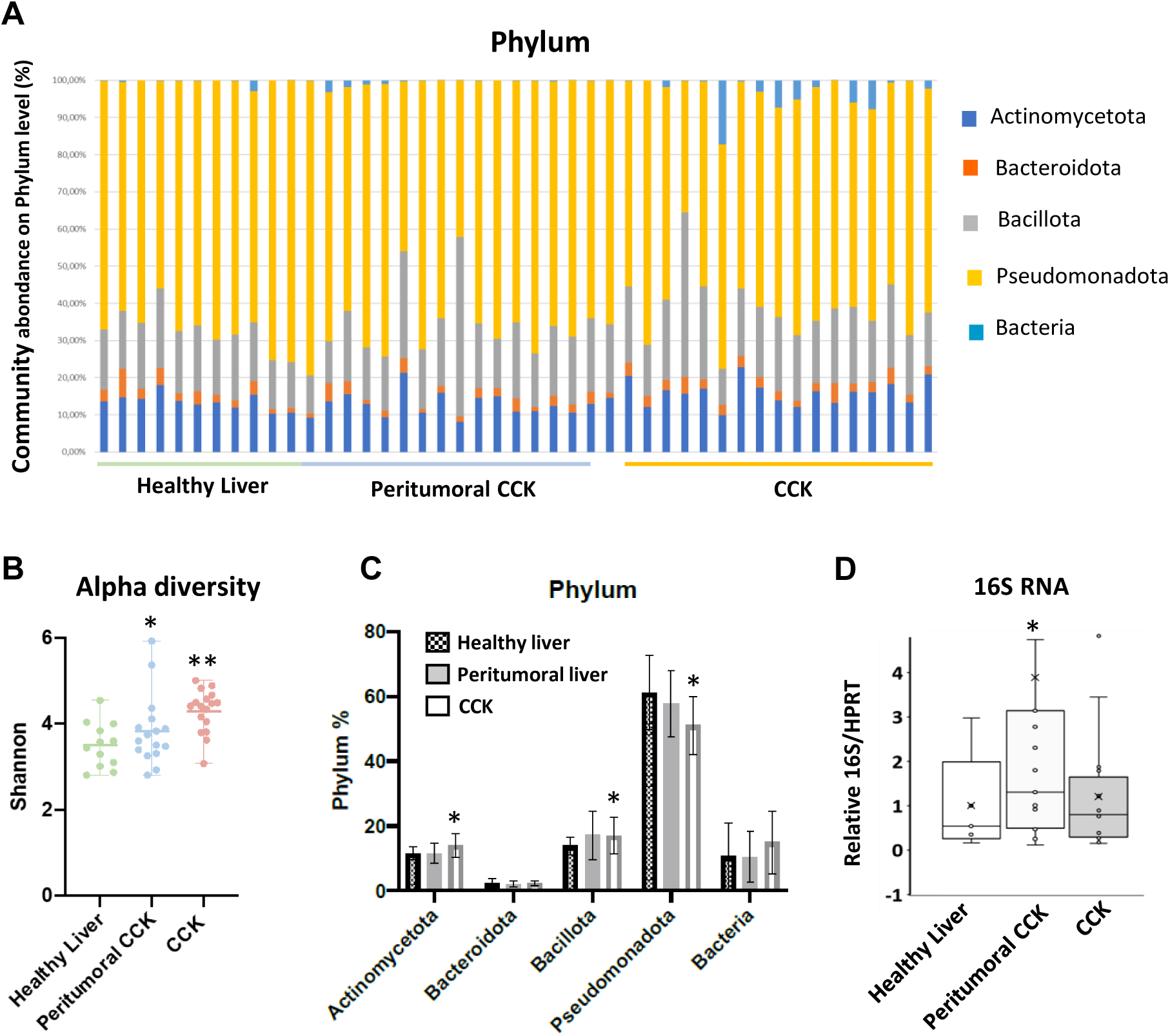
Hepatic microbiota contains bacteria expressing BamA/TamA-like proteins. A) Analysis of the composition of bacterial microbiota in healthy livers, peritumoral parenchyma and CCK. Composition features of the microbiota in each group for the 4 major phyla detected: Actinomycetota, Bacteroidota, Bacillota (Fermicutes), Pseudomonadota (Protobacteria) and undetermined Bacteria. B) Comparison of differences in the alpha diversity (Shannon index) of microbiota in the 3 groups. Statistics: *p<0.05 and **p<0.01 for alpha diversity between healthy livers versus peritumoral CCK and healthy livers versus CCK, respectively. C) Percentages of the 5 major phyla within the 3 groups of livers samples. Statistics: *p<0.05 for changes in Actinomycetota, Bacillota and Pseudomonadota healthy livers versus CCK. D) Quantification of 16S RNA by RT-pPCR in healthy and peritumoral livers and CCK using DNA primers designed to amplify ribosomal RNAs of *Providencia*, *Photorhabdus* and *Salmonella enterica* species. Statistics: *p<0.05 for significant changes between peritumoral biopsies versus healthy livers and CCK.

### 2.4. P11^Chol^ modulates the interaction of Providencia stuartii with Chol^I/T/B^-HepaRG cells

We hypothesized that Chol^I/T/B^-HepaRG cells could interact and internalize some *enterobacteriaceae* possibly via a mechanism involving the conserved peptide motif similar to P11^Chol^.To address this hypothesis, we set up a coculture model associating HepaRG cells and *Providencia stuartii* bacteria (**SI 7**) to adapt adhesion and invasion assays as previously described for *Salmonella* Heidelberg with intestinal epithelial cell cultures [58].

In these assays, Chol^I/T/B^-HepaRG cells were incubated for 30 min with *Providencia stuartii* bacteria expressing the Green Fluorescent Protein (GFP^+^) at different MOI (**Fig 4**). After washing out non-adherent *Providencia stuartii* bacteria, GFP^+^ bacteria were detected by fluorescence confocal microscopy in Chol^I/T/B^-HepaRG cell (**Fig 4A**). While some GFP^+^ bacteria were most likely bound to the cell plasma membrane since they were not detected on the same focal plane than cell nuclei (focal distance >5 μm), we observed GFP^+^ bacteria in the same focal plane than Hoechst-stained nuclear DNA (focal distance <1 μm), which strongly suggested that some bacteria were internalized (**Fig 4A**, **SI 7**). Similarly, we found some bacteria between cells by TEM and (**SI 8**). We also observed that the organization of the cytoplasm was disturbed and that these cells contained numerous large lysosome-like structures and autophagosomes containing very dense material (**SI 8**). The hypothesis of bacteria internalization was also supported by the fact that lysates of cells incubated with *Providencia stuartii* bacteria, washed and treated with gentamycin for 2 hours to kill adherent bacteria, contained viable bacteria growing and forming colonies on trypticase soy agar (**Fig 4B**). At MOI of 10, we found that the mean of bacteria adhesion was 60-fold higher than the value of internalization (**Fig 4B**).

**Figure 4.**
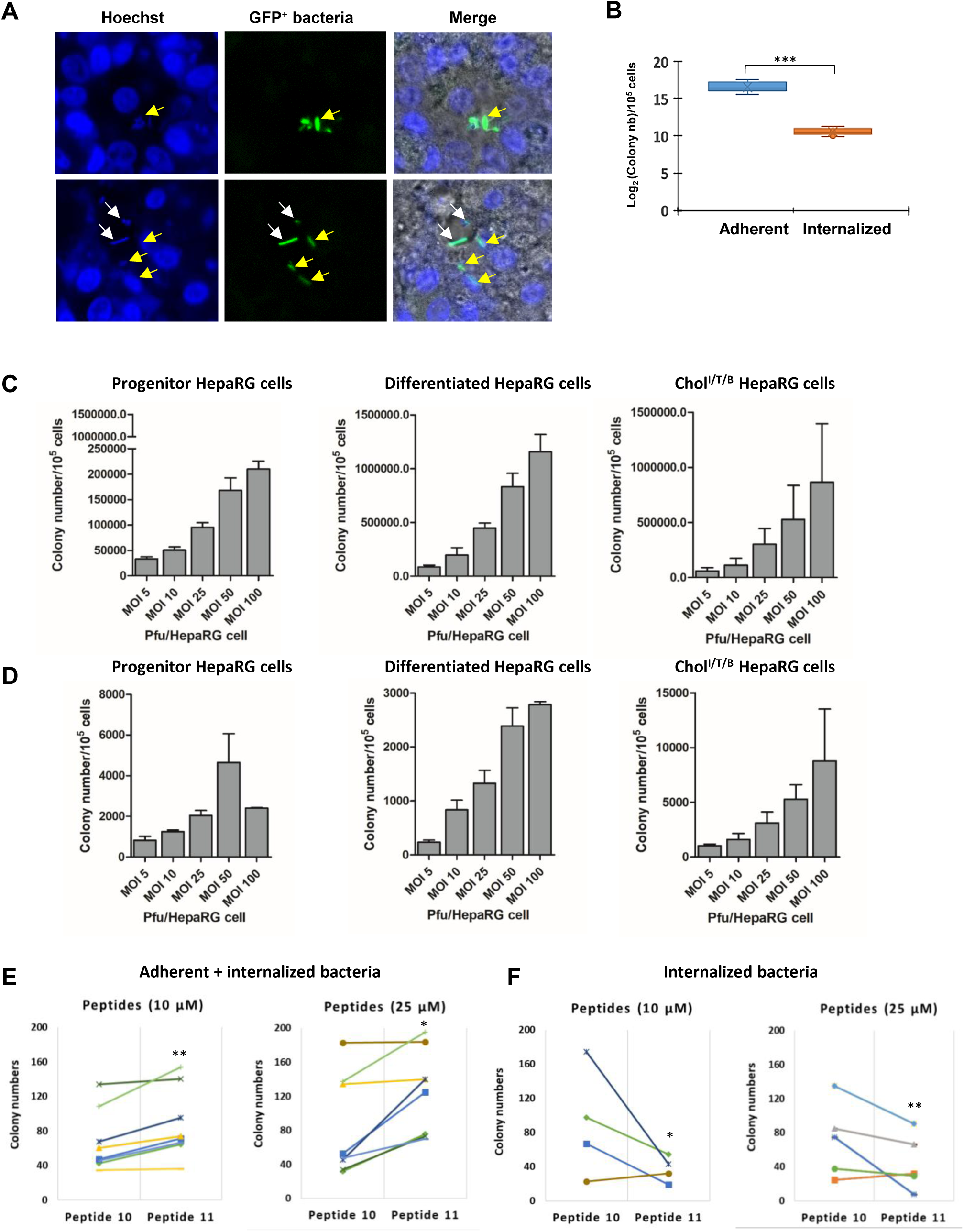
P11^Chol^ modulates internalization of *Providencia stuartii* in Chol^I/T/B^-HepaRG cells. A) Incubation of Chol^I/T/B^-HepaRG cells with GFP+ *P. stuartii* bacteria at MOI of 10 to visualize the bacteria in the human cell culture by confocal fluorescence microscopy. Bacteria are in green and nuclear DNA was stained with Hoechst. Extracellular *P. stuartii* bacteria are shown with white arrows while intracellular bacteria shown with green arrows were found on the same focal plane than nuclei in Chol^I/T/B^-HepaRG cells. B) Counts of bacteria in adhesion and internalization assays using Chol^I/T/B^-HepaRG cells infected by *Providencia stuartii* at MOI of 10. At this MOI, the means ±SD were 1.033×10^5^ ±5.09 ×10^4^ and 1.5510^+4^ ± 478 for adhesion and internalization, respectively, indicating an internalization/adhesion ratio of 1/66, *p<0.001, n=8. Quantification of adhesion (C) and internalization (D) in progenitor, differentiated and Chol^I/T/B^-HepaRG cells infected with the *P. stuartii* strain at MOI of 5, 10, 25, 50 and 100 bacteria per hepatic cell. Results were expressed in number of bacteria colonies per 10^5^ HepaRG cells. Quantification of adhesion (E) and internalization (F) in Chol^I/T/B^-HepaRG cells infected with the *Providencia stuartii* strain in the presence of peptide 10 (negative control) and P11^Chol^ at 10 and 25 μM. Statistics: *p<0.05 for significant changes in colony numbers between cultures incubated with negative control peptide 10 versus peptide P11^Chol^.

We next compared the adhesion and internalization of *Providencia stuartii* in progenitor, DMSO-differentiated and Chol^I/T/B^-HepaRG cells at different MOI (**Fig 4C, D**). Bacteria adhesion and internalization increased in a MOI-dependent manner in the HepaRG cell cultures. The adhesion was significantly higher in both Chol^DMSO^- and Chol^I/T/B^-HepaRG cells compared to that observed with progenitor cells (**Fig 4C**). The internalization values in progenitor and DMSO-differentiated cells were relatively similar and the highest colony numbers were observed in Chol^I/T/B^-HepaRG cell cultures (**Fig 4D**). In progenitor, DMSO-differentiated and Chol^I/T/B^-HepaRG cell cultures, bacterial infections at MOI 5 and 10 did not significantly affect the cell viability measured with the relative ATP content while at the higher MOI, the cell viability gradually decreased to reach 60 to 80% of the relative ATP content measured in the control cultures (**SI 8A**). At MOI 25, we observed by Transmission Electron Microscopy (TEM) dilatations of the intercellular spaces and disturbed cytoplasm organization with accumulation of large lysosome and/or autophagosome-like structures, which further reinforced the conclusion that some HepaRG cells had internalized bacteria (**SI 8B**).

We used Chol^I/T/B^-HepaRG cells infected by *Providencia stuartii* to study the effects of P11^Chol^ on bacterial adhesion and internalization (**Fig 4E, F**). At both 10 and 25 μM, P11^Chol^ increased the numbers of adherent bacteria in the Chol^I/T/B^-HepaRG cells compared to those counted in peptide 10-incubated cultures. In contrast, the amounts of internalized bacteria in the Chol^I/T/B^-HepaRG cells were reduced by P11^Chol^ compared to those found in cultures exposed to peptide 10. These data indicated that P11^Chol^ was able to modulate the interaction of *Providencia stuartii* bacteria with Chol^I/T/B^-HepaRG cells.

### 2.5. Peptides derived from Providencia stuartii and Helicobacter pylori bind to Chol^I/T/B^-HepaRG and RLEC

The biotinylated peptides corresponding to the conserved domains of BamA/TamA-like outer membrane protein from *Providencia stuartii* (peptide Provid-Stu), Pebp-like protein from *Helicobacter Pylori* (peptide Hel-Pyl), the Hepatitis C virus polyprotein (peptide HepaVC), tannase and feruloyl esterase expressed in *Alternaria alternata*, (peptide Tan-Fest) and human E-selectin (peptide E-select) were synthesized (**Table 1**) to study their binding to cholangiocytes (**Fig 5, SI 9**). More than 90% of the Chol^I/T/B^-HepaRG cells and the RLEC were labeled with peptides P11^Chol^-, Provid-Stu-, HelPyl- and E-select-Dylight^TM^488 streptavidin complexes while peptides HepaVC and Tan-Fest did not significantly bind to the HepaRG cells (**Fig 5A, B, SI 9**). When the same peptides were assayed with the HepG2 and HEK293T cells, only with E-select peptide showed a significant binding (**Fig 5C, D, SI 9**). Using peptides 10, P11^Chol^, Provid-Stu and HelPyl, no binding was found with normal human hepatocytes in primary culture, hepatic stellar-like LX2 cells and human (Huh7) and murine (Hepa1-6) HCC cell lines (**Fig 5E, SI 9**).

**Figure 5.**
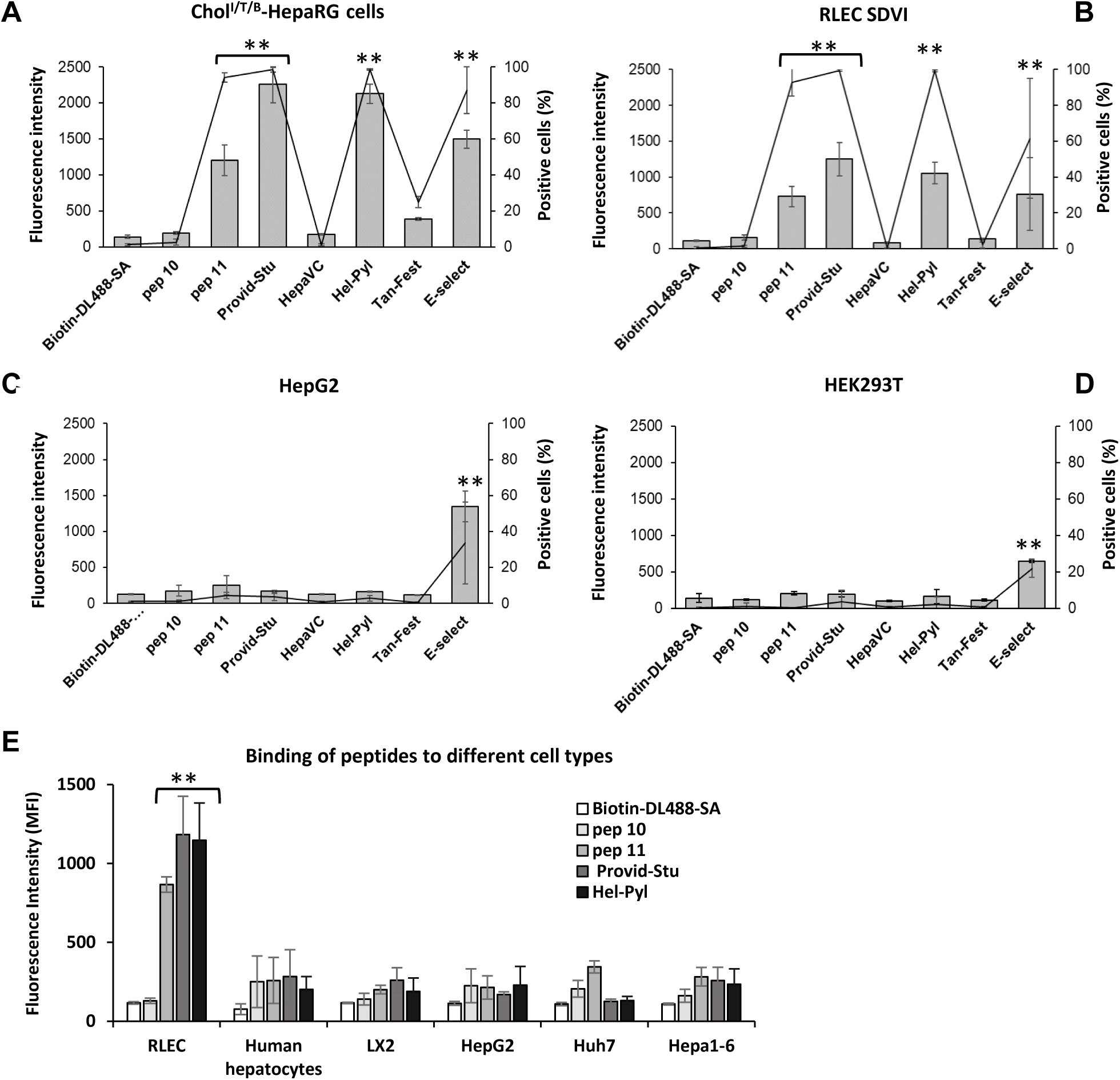
P11^Chol^ and peptides derived from *Providencia stuartii* and *Helicobacter pylori* strongly bind to Chol^I/T/B^-HepaRG and RLEC but not to normal hepatocytes and hepatocellular carcinoma cells. Cell binding assay of peptides P11^Chol^, Provid-Stu, HepaVC, Hel-Pyl, Tan-Fest and E-select-Dylight^TM^488 streptavidin complexes to Chol^I/T/B^-HepaRG cells (A), RLEC (B), HepG2 (C) and HEK2937 (D) cells expressed as relative fluorescence intensity in arbitrary units (left axis: bars) and percentage of positive cells (right axis: graphs) compared to the negative controls using peptide 10-Dylight^TM^488 streptavidin complex and Dylight^TM^488 streptavidin bound to biotin only without peptide (Biotin-DL488-SA). E) Interaction of peptides P11^Chol^, Provid-Stu, HepaVC, Hel-Pyl, Tan-Fest and E-select-Dylight^TM^488 streptavidin complexes to RLECs, normal human hepatocytes in primary culture, LX2, HepG2, Huh7 and Hepa1-6 cells. Statistics: *p<0.05 and **p<0.01 between peptide-fluorescent Dylight^TM^488 streptavidin complexes and negative controls using peptide 10-Dylight^TM^488 streptavidin complex and Dylight^TM^488 streptavidin bound to biotin.

### 2.6. Detection and isolation of rat and human hepatic cholangiocytes using P11^Chol^

Then, we examined whether peptides P11^Chol^-, Provid-Stu- and HelPyl-Dylight^TM^488 streptavidin complexes could be used as fluorescent ligands to detect and isolate intrahepatic liver (LEC) and/or biliary epithelial (BEC) cells by flow cytometry from cell suspensions obtained after dissociation of rat and human liver parenchyma (**Fig 6A, B**). Dot-plots [size (FSC) versus fluorescence (FITC)] obtained for rat and human hepatic cell suspensions incubated with peptide 10- and P11^Chol^-Dylight^TM^488 streptavidin complexes evidenced 4.1%±1.3 and 3.9%±1.7 of P11^Chol^-positive cells (**Fig 6A, B**) in rat and human cells, respectively. In contrast, no positive cells were detected with Provid-Stu- and HelPyl-Dylight^TM^488 streptavidin complexes.

**Figure 6.**
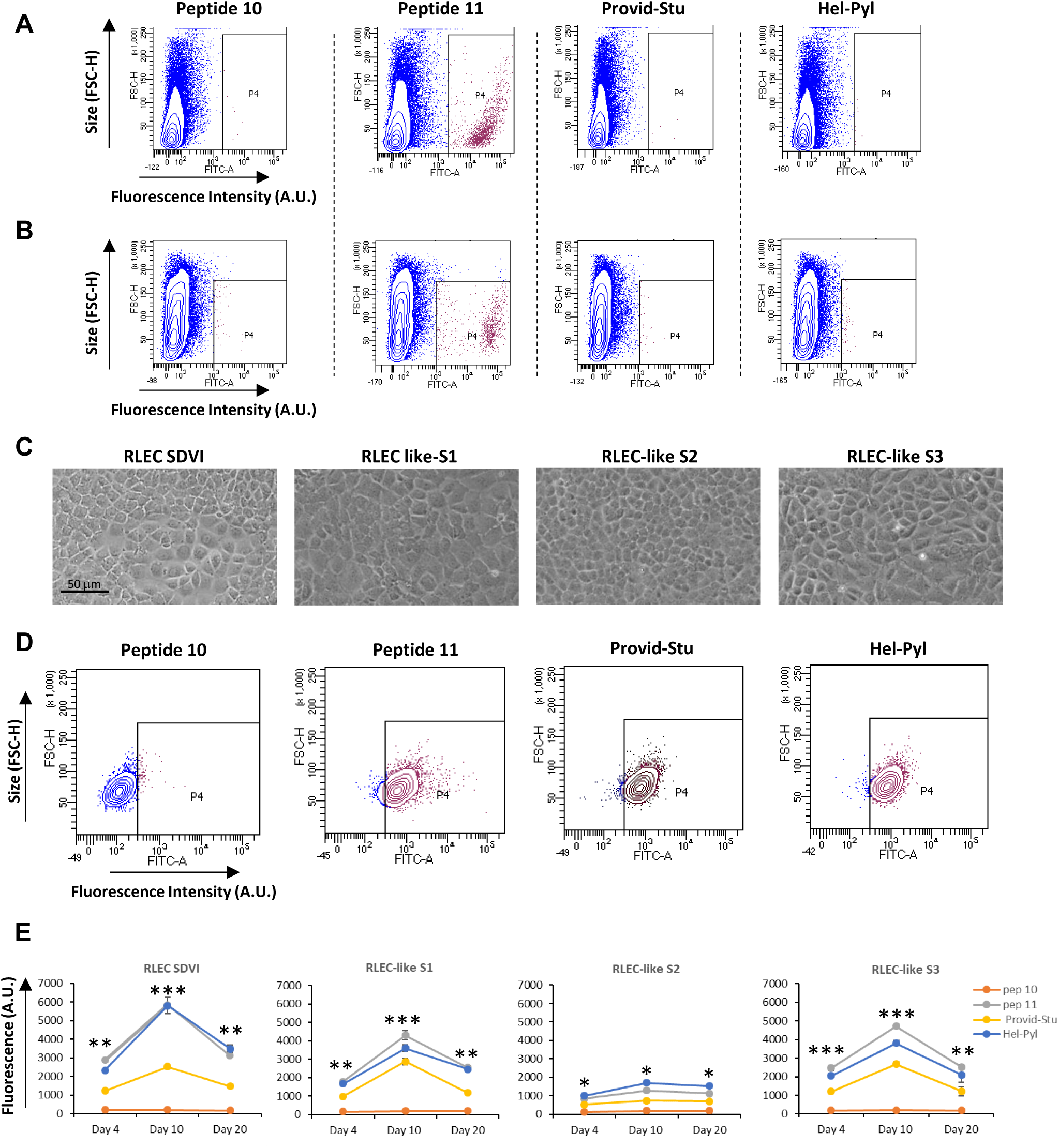
Detection and of rat and human hepatic P11^Chol^ positive cells in total liver cell suspensions and isolation of RLEC-like cells. Cell binding assay of peptides 10, P11^Chol^, Provid-Stu and Hel-Pyl-Dylight^TM^488 streptavidin complexes to rat (A) and human (B) whole liver cell suspensions following enzymatic dissociation of the liver parenchyma. The negative population was defined using the peptide 10-Dylight^TM^488 streptavidin complex. Positive population was detected and quantified in P4 window following incubation of hepatic cells with P11^Chol^-Dylight^TM^488 streptavidin complexes while no positive cells were detected with Provid-Stu and Hel-Pyl-Dylight^TM^488 streptavidin complexes. C) P11^Chol^-Dylight^TM^488 streptavidin complex positive hepatic cells from 3 different rat livers were sorted by FACS and expanded to establish stable RLEC-like S1, S2 and S3 cell lines with cell morphologies similar to RLEC SDVI cells. After establishing stable cell lines (passage 2), RLEC-like S1 cells (D) as well as RLEC SDVI, RLEC-like S2 and S3 cells (E) became positive for binding to peptides P11^Chol^, Provid-Stu and Hel-Pyl-Dylight^TM^488 streptavidin complexes. Statistics: *p<0.05, **p<0.01 and ***p<0.001 between peptides P11^Chol^-, Provid-Stu- and Hel-Pyl- and peptide 10-Dylight^TM^488 streptavidin complexes.

We sorted P11^Chol^ positive cells by FACS from 3 distinct liver suspensions and obtained stable lines further referred as RLEC-like S1, 2 and 3 cell lines (**Fig 6C**). These epithelial cells proliferate at low density to form confluent squamous monolayers of polygonal cells (**SI 10**), which can be expanded over many passages demonstrating their potential of proliferation *in vitro*. RLEC-like S1 and 3 cells showed very similar morphology compared to the previously described RLEC SDVI cell line [55] while RLEC-like S2 cells appeared smaller compared to the other RLEC cells (**Fig 6C, SI 10**). Unexpectedly, we found that the 3 peptides P11^Chol^-, Provid-Stu- and HelPyl-Dylight^TM^488 streptavidin complexes bound to the RLEC-like S1, S2 and 3 cells (**Fig 6D, E**) as observed for RLEC SDVI cells although no positive cells were detected with Provid-Stu- and HelPyl peptides in the liver cell suspension used to establish RLEC-like cell lines (**Fig 6A**). In addition, we found that the intensities of fluorescence were relatively similar for RLEC SDVI, RLEC-like S1 and 3 cells incubated with peptides P11^Chol^- and HelPyl-while fluorescence levels were lower with Provid-Stu peptide in all cell lines. Moreover, the fluorescence means fluctuated in proliferating versus quiescent cells at different days after plating with maximal intensities at day 10 (**Fig 6E**). In RLEC-like S2 cells, the binding of peptides P11^Chol^, Provid-Stu and HelPyl streptavidin complexes was much weaker compared that observed with the other epithelial cells.

### 2.7. Characterization of rat liver epithelial cell lines

In order to characterize RLEC S1, S2 and S3 cells, cultures were maintained confluent and cell morphologies were monitored for 60 days (**Fig 7A, SI 10**). After 2 to 3 weeks, high density cell monolayers showed squamous appearance and abundant deposition of extracellular matrix fibers. In addition, some cells were organized into spherical and cyst-like structures adhering to the cell monolayer or floating in the culture medium (**SI 10**). Using TEM, we observed abundant extracellular matrix (ECM) components organized in large fibers deposited between cells (**Fig 7A**). Ultrastructure of RLEC cells also evidenced microvilli and very abundant mitochondria. The production of ECM was confirmed by RT-qPCR, which showed that collagen 1A1 mRNA levels were very low in proliferation RLEC cells 4 days after plating but were strongly enhanced in quiescent cells at days 10 and 20 (**Fig 7B**). We also demonstrated that confluent quiescent RLEC SDVI, S1, S2 and S3 cells expressed cholangiocyte cell membrane markers such as CD24 and CFTR while CD133 expression was either very low or undetectable (**Fig 7C, SI 11**).

**Figure 7.**
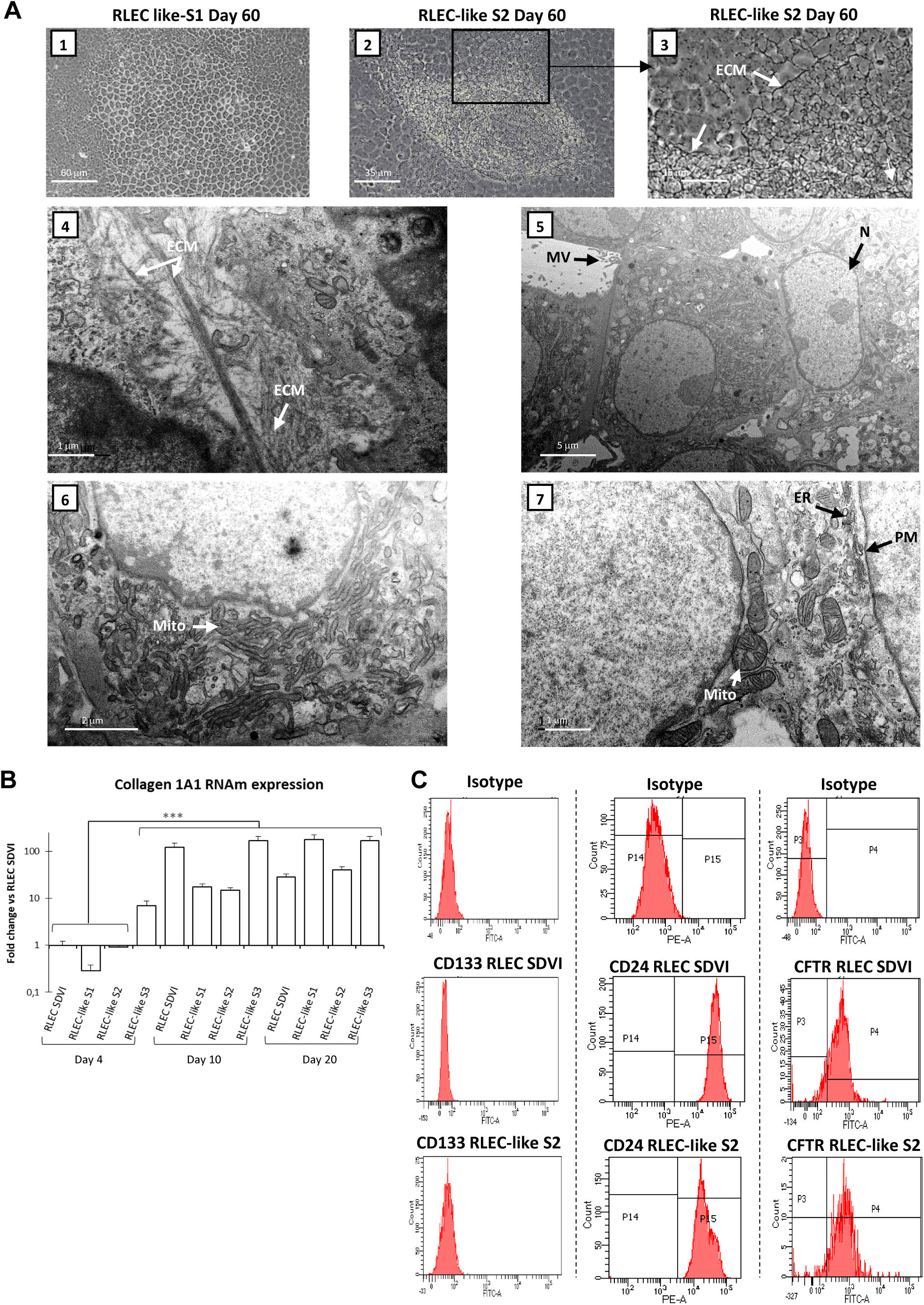
Morphology and ultrastructure of CD24 and CFTR positive RLEC-like cells. Cell morphology in phase contrast microscopy (A1-3) and ultrastructure (A4-7) in transmission electronic microscopy (TEM) of RLEC-like cells was studied after establishing stable RLEC-like S1, S2 and S3 cell lines. Confluent squamous monolayers of RLEC-like S1 and S2 cells (A1-3) characterized by deposition of fibers of extracellular matrix (ECM) confirmed by TEM (A4). Ultrastructure of RLEC-like cells, illustrated with photographs of RLEC-like S2 cells (A4-7) was characterized by polygonal cell shape with plasma membrane (PM) presenting microvilli (MV), very abundant mitochondria (Mito) and endoplasmic reticulum (ER) and a single nucleus (N) with one or two nucleoli. B) Collagen 1A1 mRNA RT-qPCR in proliferating RLEC cells 4 days after plating and in quiescent cells at days 10 and 20 after seeding. C) Flow cytometry analysis in RLEC SDVI and RLEC-like S2 cells for the detection of CD133, CD24 and CFTR expression compared to background fluorescence signal obtained with control isotype antibodies.

We next studied whether RLEC could self-organize on Matrigel matrix to form organoids, three-dimensional cysts or branching tubular structures as previously described for pluripotent stem cell-derived cholangiocytes [59, 60] and Chol^I/T/B^-HepaRG cells [52]. RLEC SDVI, S1, S2 and S3 cells were plated either on collagen I or Matrigel coated dishes and cell morphologies were monitored for 10 to 18 days (**Fig 8A**). When plated on collagen I, RLEC cells formed squamous monolayers as observed on plastic support. In contrast, RLEC cells seeded on Matrigel did not spread and formed small cell clusters, which evolved towards complex multicellular organoids some of them exhibiting branched tubular structures (**Fig 8A**).

**Figure 8.**
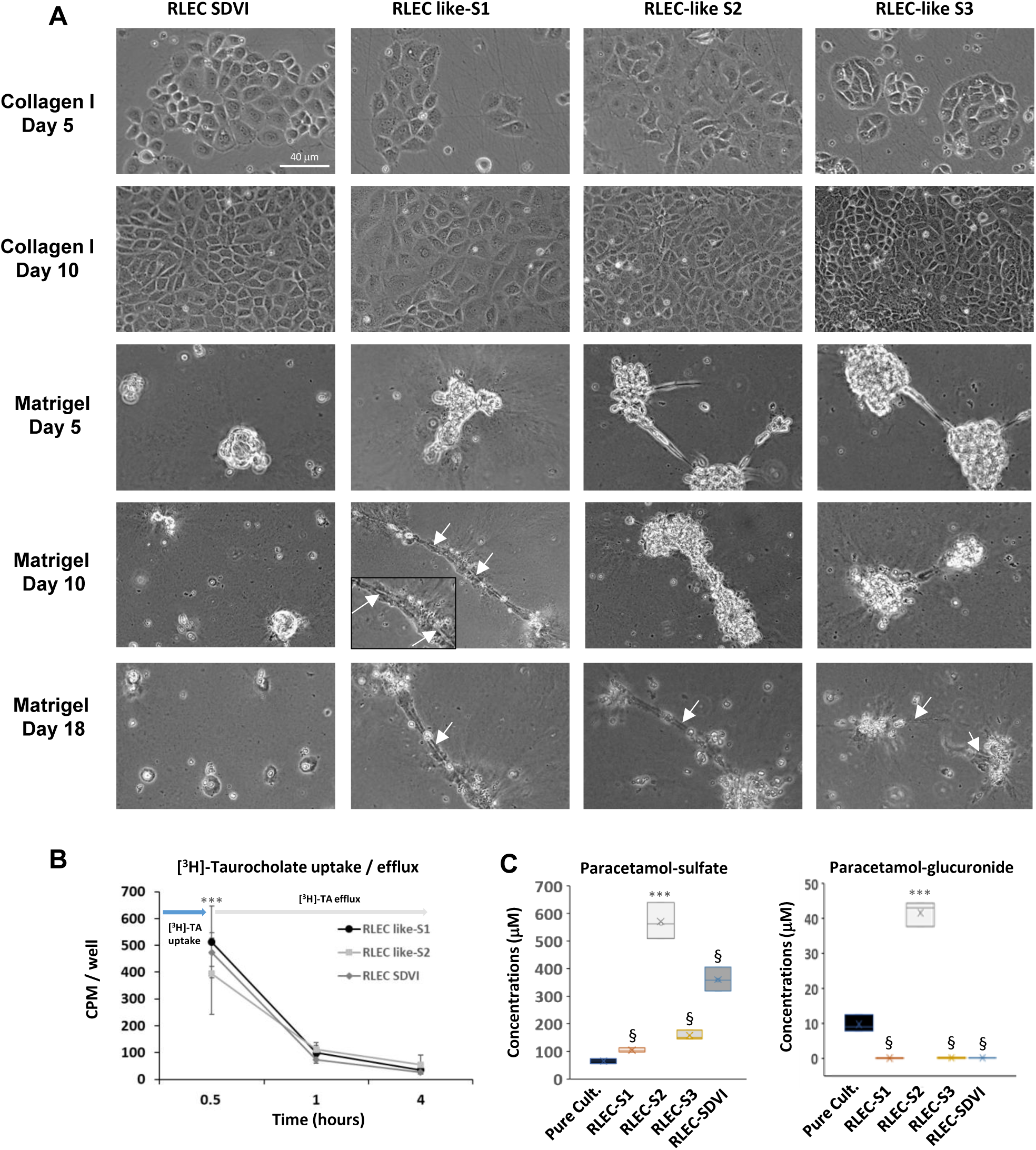
RLEC-like cells form spheroids on Matrigel coating and express functional cholangiocyte markers. A) Morphology in phase contrast microscopy of RLEC SDVI and RLEC-like S1, S2 and S3 cells at days 5, 10 and 18 after plating on collagen I and Matrigel coating. RLEC cells form organoids, three-dimensional cysts or tubular structures (white arrows) on Matrigel matrix. B) Uptake measured during 30 min (0.5h) and efflux (clearance at 1 and 4 hours) of [^3^H]-taurocholic acid (75 MBq/well) in confluent RLEC SDVI and RLEC-like S1 and S2 cell lines. C) Analysis by liquid chromatography coupled to tandem Mass Spectrometry (LC/MS-MS) of sulfotransferase and uridine 5′-diphospho-glucuronosyltransferase enzymatic activities by measuring the production of sulfate- and glucuronide-paracetamol conjugates in confluent RLEC SDVI, S1, S2 and S3 cells compared to pure cultures of normal rat hepatocytes. Statistics: §p<0.05, between RLEC SDVI, S1 and S3 cells versus pure cultures of hepatocytes (Pure Cult.) and ***p<0.001 between RLEC-like S2 cells and all the other conditions.

Since cholangiocytes are involved in metabolism and transport of bile salts, we also studied the uptake and efflux of radiolabeled taurocholate in confluent RLEC cells (**Fig 8B**). RLEC cells accumulated significant amounts of [^3^H]-taurocholate demonstrating their ability to import this bile acid. After washing out the [^3^H]-taurocholate present in the culture medium, we showed that the intracellular [^3^H]-taurocholate contents rapidly decreased after 1 and 4 hours demonstrating its active efflux from RLEC cells.

In the liver, phase I and II drug metabolism enzymes are mainly expressed in hepatocytes, however, some sulphotransferases have been shown to be expressed and active in rat and human cholangiocytes [61]. We first studied whether RLEC cells catalyzed phase I drug biotransformation using dextromethorphan, bupropion and midazolam, which are oxidized by CYP2D6, CYP2B6 and CYP3A4/5 enzymes, respectively. As expected, we did not find any metabolites of these substrates in RLEC cell media (data not shown). In contrast, we evidenced that all RLEC cells produced sulfate-paracetamol conjugates (**Fig 8C**). The RLEC-like S2 cells showed much higher paracetamol sulfonation activity compared to other RLEC cells and rat hepatocytes in pure cultures. Unexpectedly, we also found that RLEC-like S2 cells produced glucuronide-paracetamol conjugates at higher levels than normal rat hepatocytes (**Fig 8C**) indicating that these cells expressed uridine 5′-diphospho-glucuronosyltransferases (UGT).

### 2.8. RLEC allow long-term survival of normal rat hepatocytes in coculture

We next determined whether RLEC-like S1, S2 and S3 cells were able to promote long-term survival and expression of liver specific functions of hepatocytes in coculture associating these two cell types as previously described for RLEC SDVI cells [55]. After plating, isolated rat hepatocytes rapidly attached to culture dishes to form colonies of cuboidal cells showing dense cytoplasm, large nucleus with single nucleolus and bile neo-canaculi (**SI 12**). Twenty-four hours after plating hepatocytes, RLEC cells were seeded and they attached to culture dishes between hepatocyte colonies to form confluent cocultures 2 days later (**Fig 9A, SI 12**). As expected, the hepatocytes lost their typical morphology after 3 days in pure culture. Their cytoplasm became clearer, the nuclei enlarged and showed fragmented nucleoli. After 3 to 5 days, some cells detached characterizing progressive cell death by apoptosis. In cocultures with RLEC-like and SDVI cells, hepatocytes kept their cuboidal shape, exhibited bile canaliculi and maintained well-defined colonies surrounded by RLEC cells (**Fig 9A, SI 12**). We confirmed that apoptosis took place between days 3 and 7 in pure culture of hepatocytes by showing the strong increase in caspase activity while apoptosis remained very low in cocultures confirming the long-term survival of hepatocytes associated with RLEC cells (**Fig 9B**). In addition, at days 7 and 14, albumin mRNA levels were significantly higher in cocultures than in pure cultures (**Fig 9C**). Similarly, midazolam 1’-hydroxylase (CYP3A4/5), bupropion hydroxylase (CYP2B6), dextrometorphan O-demethylase (CYP2D6) and paracetamol sulfonation and glucuronidation activities were 2 to 10-fold higher in cocultures than in pure cultures (**Fig 9D**).

**Figure 9.**
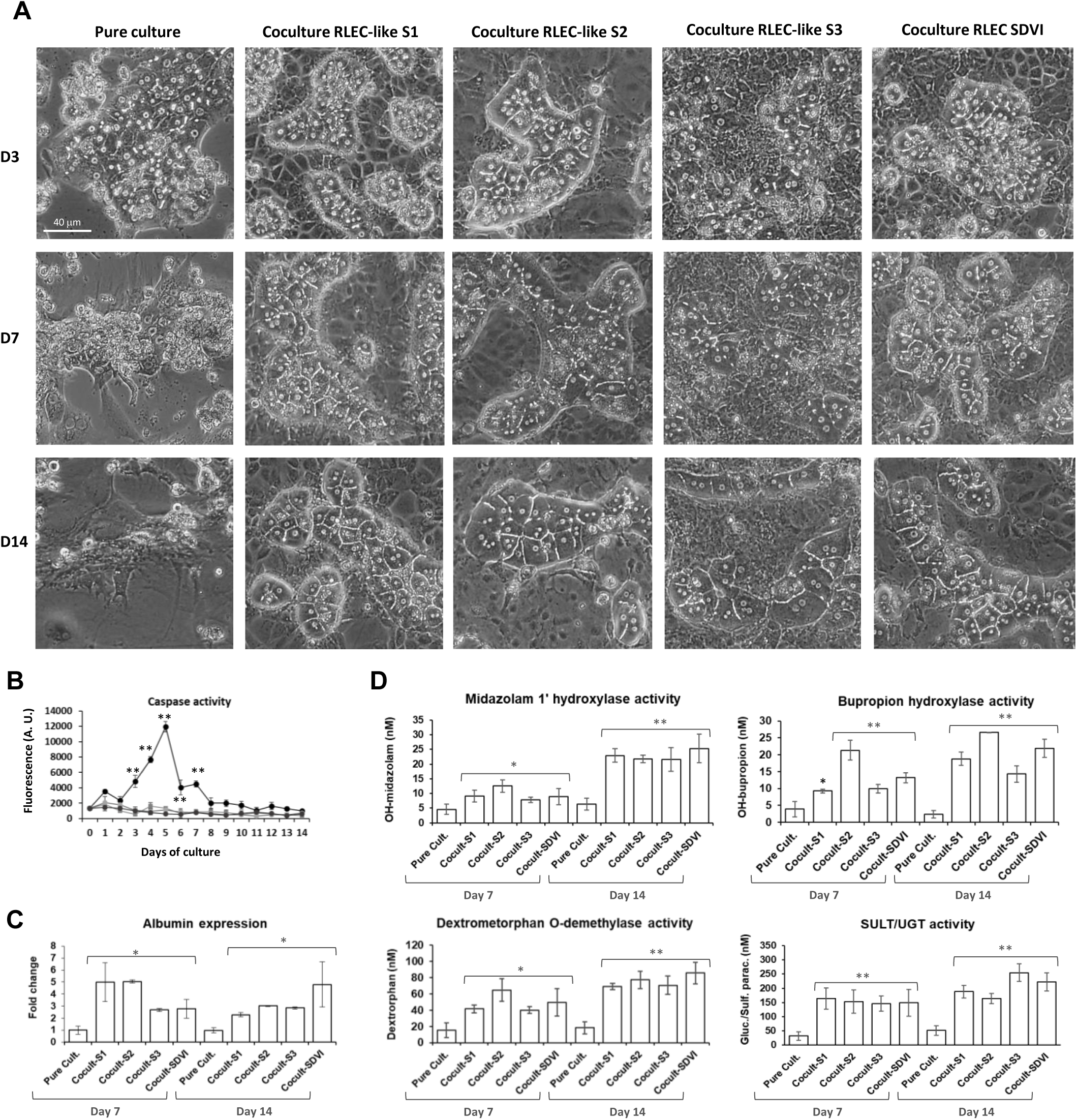
RLEC cells promote long-term survival of normal rat hepatocytes in coculture. A) Morphology in phase contrast microscopy of normal rat hepatocytes in pure cultures and cocultures with RLEC SDVI and RLEC-like S1, S2 and S3 cells at days 3, 7 and 14 after plating. B) DEVD-AMC caspase activities (fluorescence) measured in normal rat hepatocytes pure cultures (dark circles) and cocultures (grey circles and squares) with RLEC SDVI and RLEC-like S1 and S2 cells between days 1 and 14 after plating. Albumin mRNA RT-qPCR (C) and midazolam 1’-hydroxylase (CYP3A4/5), bupropion hydroxylase (CYP2B6), dextrometorphan O-demethylase (CYP2D6) and paracetamol sulfonation and glucuronidation activities (D) in normal rat hepatocytes pure cultures (Pure Cult.) and cocultures (Cocult.) prepared with RLEC SDVI and RLEC-like S1, S2 and S3 cells at days 7 and 14 after plating. Statistics: *p<0.05 and **p<0.01 between pure cultures versus cocultures.

The coculture associating hepatocytes and RLEC cells is also characterized by branched bile canaliculus network between hepatocytes, which is stable for several weeks while bile canaliculi visible in pure cultures of hepatocytes during 3 to 5 days disappeared thereafter (**Fig 9A, SI 12**). We performed a semi-quantitative analysis of the bile canaliculi network at day 7 after plating in pure cultures and cocultures with RLEC-like S1, S2, S3 and SDVI cells by detecting cholyl-lysyl-fluorescein (CLF) efflux (**Fig 10, SI 13**). As expected, very few bile canaliculi were detected in pure cultures of hepatocytes (**Fig 10A**). In contrast, CLF accumulation delineated functional bile canaliculi forming branched network in hepatocyte colonies in RLEC-like S1, S3 and SDVI cells (**Fig 10A**, **Fig 10B**) while cocultures prepared with RLEC-like S2 cells exhibited a far less intense CLF staining compared to the bright networks observed with the other cocultures, although dilated canaliculi could be observed by phase contrast microscopy in cocultures with RLEC-like S2 cells (**Fig 9A**, **10A**).

**Figure 10.**
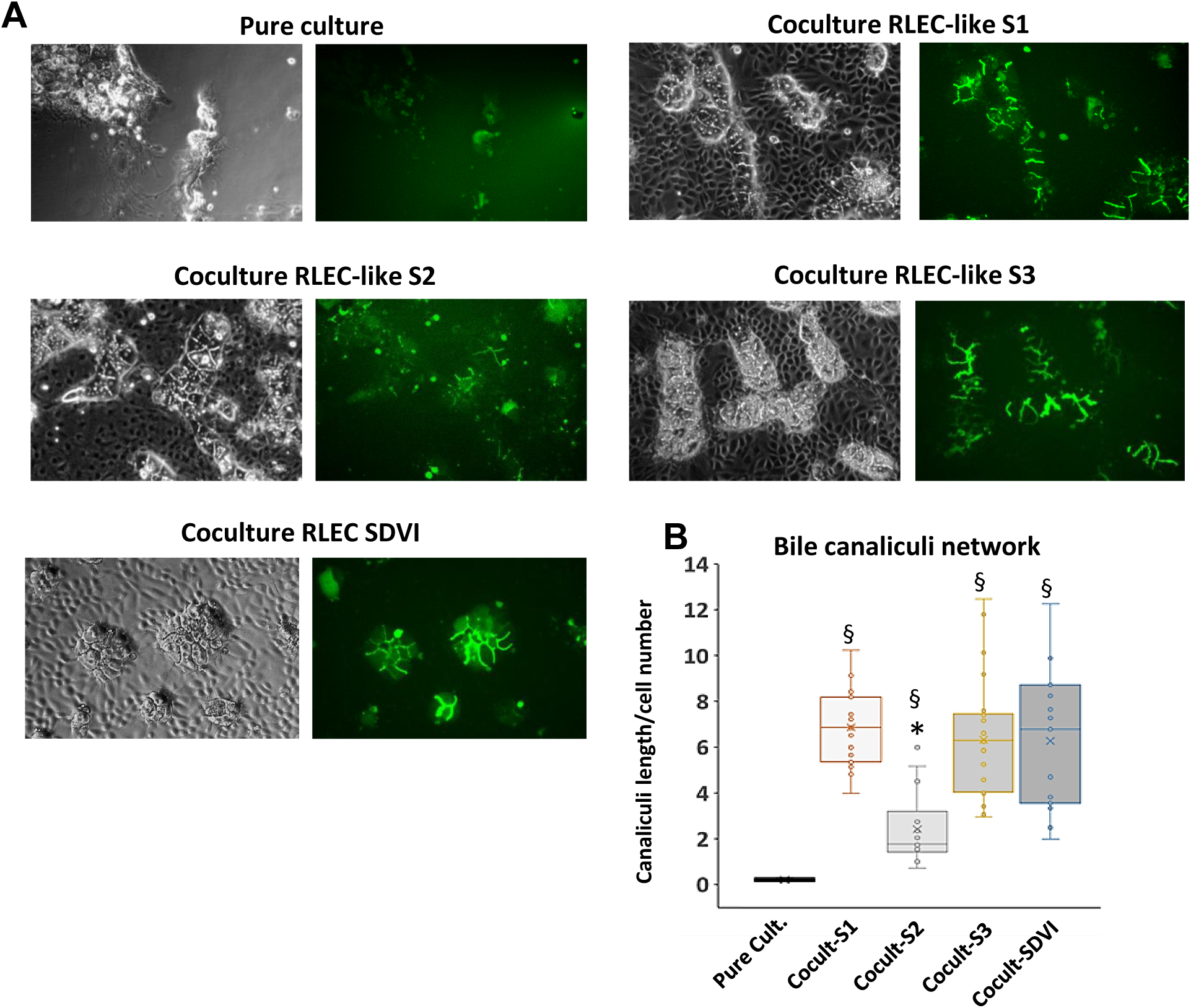
Analysis of functional bile canaliculi in hepatocytes cocultured with RLEC cells. Accumulation of cholyl-lysyl-fluorescein (CLFI) detected in fluorescence microcopy (A) was used to quantify of the canaliculus network in rat hepatocyte colonies cocultured with RLEC-SDVI as well as RLEC-like S1, S2 and S3 cells (B). Length of bile canaliculi were measured for colonies including at least 6 hepatocytes or more. Hepatocytes were counted in colonies and ratio of canaliculi/number of hepatocytes was established. Results were obtained from 4 independent experiments in which at least 6 images (fields of cocultures and a total of 250 to 500 hepatocytes) were analyzed. Statistics: §p<0.05, between pure cultures of hepatocytes (Pure Cult.) and coculture of hepatocytes and RLEC cells (Cocult.), *p<0.05 between cocultures prepared with RLEC-like S2 cells (Cocult-S2) versus the other cocultures.

## 3. Discussion

In this study, we identified the peptide P11^Chol^ [seq: TFLNSVPTYSYWGGGS] that binds to human and rat cholangiocytes and liver epithelial cells. This peptide shows strong similarities with a conserved peptide motif [seq: L-N-(S/N)-(V/I)-P-T-Y-Y-W-G-X-G] found in BamA/TamA-like outer membrane proteins expressed in *enterobacteriaceae* belonging to Pseudomonadota phylum. In gram-negative bacteria, β-barrel BamA-B-C-D-E assembly machinery complexes are inserted into the outer membrane by the conserved ∼85 kDa OMP85/BamA proteins containing a C-terminal 16 β-stranded barrel structure interacting with the other subunits [762, 63]. This complex and the OMP85/BamA barrels show remarkable conserved structural features ensuring the functional interaction of the different subunits to transport small molecules [62, 63]. Using crystal structures of different TamA-like proteins, we found that the consensus peptide motif was located in different positions of the OMP85/BamA protein barrel structures depending on bacteria species. This peptide motif is located within the barrel of BamA/TamA-like proteins for *Escherichia coli* and *Salmonella enterica* and in a short extracellular loop in *Providencia stuartii*’s protein. This peptide motif is thus most likely exposed on the external face in some *enterobacteriaceae* although BamA/TamAl proteins are subjected to conformational changes during transporter activity [62, 63].

We also evidenced the presence of *enterobacteriaceae* in healthy human livers, CCK and peritumoral livers as previously reported [64–66] and showed significant quantitative variations in the abundance in Actinomycetota, Bacillota and Pseudomonadota between healthy livers and CCK. Recently, it was demonstrated that normal and pathological livers hosted microbiota that may play important role in hepatic immunity [64] and during progression of hepatocellular-[65] and cholangio-carcinomas [66]. The presence of bacteria in the vicinity of intrahepatic cholangiocytes could result from gut inflammation, dysbiosis and translocation of pathogens across gut epithelial cells [67]. In addition, commensal bacteria and pathogens such as *Salmonella* strains are able to colonize the gallbladder of patients with cholecystitis [68] and bile components regulate virulence gene expression and stress responses [69].

While many articles described the internalization of various bacteria species by human cells [58, 68, 70], we did not find in the literature any report describing an *in vitro* model of cholangiocytes to assess *enterobacteriaceae* adhesion and/or internalization. Herein, we report that *Providencia stuartii* bacteria bind to cholangiocyte-like HepaRG cells based on confocal microscopy and TEM data and quantification of bacteria colonies from cell extracts supporting the conclusion that *Providencia stuartii* can also be internalized in these cells. In this model, we also demonstrated that peptide P11^Chol^ increased the adherence of *Providencia stuartii* bacteria onto Chol^I/T/B^-HepaRG cells while reducing the internalization, which supports the hypothesis that binding of *enterobacteriaceae* to cholangiocytes could involve Tam/BamA-like proteins and the conserved peptide motif [L-N-(S/N)-(V/I)-P-T-Y-Y-W-G-X-G].

In a second part of this work, we demonstrated that the fluorescent peptide P11^Chol^ is suitable to detect and isolate rat and human LEC by flow cytometry from whole hepatic cell suspensions obtained after dissociation of liver parenchyma. We isolated by cell sorting rat liver epithelial P11^Chol^-positive cells and established three distinct RLEC-like cell lines, which are morphologically very similar to RLEC SDVI previously isolated by cell cloning in our laboratory [55]. We showed that the newly established RLEC-like and RLEC SDVI cells bind the peptide P11^Chol^ as well as the peptides Provid-Stu and Hel-Pyl present in BamA/TamA-like outer membrane proteins of *Helicobacter Pylori* and *Providencia stuartii*, respectively. The binding of the three peptides to all RLEC cell lines was unexpected since only peptide P11^Chol^ and not peptides Provid-Stu and Hel-Pyl bound to LEC from the whole liver cell population. These data suggest that the three peptides may bind to different proteins and/or that phenotypical changes of LEC occurring *in vitro*.

In adult liver, the intrahepatic biliary tree contains two distinct types of liver epithelial cells (LEC) often referred as small and large cholangiocytes preferentially located in bile ductules and large hepatic ducts, respectively [59, 60]. Small cholangiocytes are able to proliferate *in vivo* while large ones are mostly quiescent [71]. Both small and large cholangiocytes express EpCAM, CD24 and CD133[59, 60], but some proteins are expressed only in large cholangiocytes such as CFTR, the somatostatin receptor 2 and the secretin receptor [60].

For many years, isolation of LEC was a complex and time-consuming procedure including multiple steps of cell cloning to obtain pure LEC/BEC populations from normal rat and mouse livers [71–76] or following ductular proliferation obtained by bile duct ligation in rats [74, 77] and expanded *in vitro* [78]. Recently, human LEC populations were sorted from whole liver cell suspension by selecting the cells that were CD45^-^/CD31^-^ but expressing EpCAM and CD24. In this population, CD133^+^ or CD133^-^ delineated two sub-populations of LEC with a progenitor-like profile for CD133^+^ cells [79, 80]. Both murine and human LEC can generate functional cholangiocytes *in vitro* under appropriate culture conditions [73, 75, 81, 82] and form bile duct structures in mice [79].

The present study provides new means to isolate primary liver epithelial cells. We demonstrated that peptide P11^Chol^ labeled ∼4% of cells within the whole hepatic cell population, in agreement with the 5% of cholangiocytes estimated among the liver cells [60]. The RLEC cell lines isolated and characterized in our study express CD24 but were negative for CD133. RLEC-like S1/S3 and RLEC SDVI cells express the CFTR channel, a well-established cholangiocyte marker [60, 79]. RLEC-like S2 cells also express CFTR but at a lower level than the other cell lines. We further characterized these RLEC cell lines and showed that they 1) produce a very abundant extracellular matrix, 2) form spheroids and tubular structures when plated on Matrigel, 3) actively transport bile salts, 4) catalyze paracetamol sulfonation, especially RLEC-like S2 cells demonstrating that they express one or several SULTs involved in paracetamol sulfonation [83], 5) support long-term survival of rat hepatocytes, the expression of various liver/hepatocyte specific functions and the development of functional canaliculi in cocultures.

Our data also supported the conclusion that RLEC-like S2 cells are phenotypically different from their S1 and S3 counterparts and RLEC SDVI since RLEC-like S2 cells show a different cell morphology, a weaker binding to peptide P11^Chol^ and a lower CFTR expression. In addition, they exhibit high SULT and produce high amounts of glucuronide paracetamol conjugates while other RLEC do not, indicating that they express UGT1A1, UGT1A6, and/or UGT1A9, the major UGTS producing paracetamol glucuronides [84]. In addition, RLEC-like S2 cells do not promote the expansion of bile canaliculi in hepatocytes. These phenotypical differences support the concept of small and large cholangiocytes and the general consensus on variations in differentiation between cholangiocytes within the Hering ducts in contact with the hepatocytes and those forming the bile ducts of the portal space. Although these different cholangiocytes express common biliary markers such as CD24 and EpCAM, they also express specific proteins characterizing these differentiation states such as the CFTR found in large but not in small cholangiocytes [59, 60]. In agreement, RLEC-like S2 cells that are smaller cells than the other cell lines show a weaker expression of CFTR. In this context, the quantitative differences in the binding of peptide P11^Chol^ to RLEC cell lines and the SULT and UGT enzymatic activities may constitute new markers that further distinguish the different intrahepatic cholangiocytes.

## 4. Materials and methods

### 4.1. Cell cultures and bile canaliculus staining

Human HepaRG cells were cultured as previously described [51, 54, 85] at 37°C and 5% CO_2_ atmosphere in William’s E medium (Gibco, Thermo Fisher Scientific, Waltham MA, USA) supplemented with 10% Fetal calf serum (mix 1/3: Biosera, FB-1001/500, HyClone Fœtal II SH30066.03), 100 units/mL penicillin, 100µg/mL streptomycin, 2mM L-glutamine, 5μg/mL insulin and 10^-5^M sodium hemisuccinate hydrocortisone (Upjohn Pharmacia, Pfizer, New York USA). Progenitor HepaRG cells seeded at low density (2.6×10^4^ cells/cm^2^) (**SI 1A**) actively proliferated during the first week (**SI 1B**). Two weeks after plating, the differentiation of committed hepatocytic and biliary cells (**SI 1C**) was potentiated by culturing the cells for 2 more weeks in medium supplemented by 2% of DMSO (**SI 1D**) to obtain differentiated hepatocyte-like and cholangiocyte-like HepaRG cells [51]. Hepatocyte- and cholangiocytes-like HepaRG cells were selectively detached using mild trypsinization with trypsin 0.05% diluted in phosphate-buffered saline (PBS) (vol/vol) as previously described [51] and plated separately at high density (2.5×10^5^ cells/cm^2^) to obtain cultures enriched in hepatocytes or cholangiocytes (**SI 1E-F**). Both hepatocyte- and cholangiocyte-like cells were cultured in medium supplemented with 2% DMSO. These cholangiocyte-like cells were referred as to Chol^DMSO^-HepaRG cells. Human hepatocytes (PHH) were seeded at a density of 2.5×10^5^ cells/cm^2^ and cultured in same medium than differentiated HepaRG hepatocytes.

Cholangiocyte-like cells were also obtained through a second procedure: progenitor HepaRG cells at low density were treated for 2 days with Interleukin-6 (10 ng/mL, Miltenyi Biotec SAS, Paris, France), then for 2 days with sodium taurocholate hydrate (10nM) and for 2 more days with sodium taurocholate hydrate (10nM) plus sodium butyrate (1.8μM). These cholangiocyte-like cells were referred as to Chol^I/T/B^-HepaRG cells [52].

HepG2 cells, obtained from American Tissue Culture Collection (ATCC, Los Altos, CA, USA) were seeded at a density of 6.6×10^4^ cells/cm^2^ in MEM Eagle medium supplemented with 10% FBS, 50 UI/mL penicillin, 50µg/mL streptomycin and 2mM L-glutamine. The HuGB cells were maintained in Williams’ E medium containing antibiotics and 10% FCS as previously described [56]. LX-2 human hepatic stellate cells (Sigma-Aldrich, St. Louis MO, USA) were maintained in DMEM medium supplemented with 2% FBS, 5UI/mL penicillin, 50µg/mL streptomycin and mM L-glutamine and were used at low passage numbers. The human bronchial carcinoid NCI-H727 cells and the human embryonic kidney 293 (HEK293T) cells were respectively cultured in RPMI 1640 and DMEM media (Gibco, Thermo Fisher Scientific, Waltham MA, USA) supplemented with 10% FCS 50 UI/mL penicillin, 50µg/mL streptomycin and 2mM L-glutamine.

Rat liver epithelial cell (RLEC) line SDVI was established as previously described [76] from livers of 10-day old Sprague-Dawley rats in our laboratory [55]. RLEC SDVI cells were cultured in William’s E medium supplemented with 10% Fetal calf serum (HyClone Fœtal II SH30066.03), 100 units/mL penicillin, 100µg/mL streptomycin, 2mM L-glutamine and were used between passages 10 and 30.

Hepatocytes and non-parenchymal liver cells were isolated by a two-step collagenase perfusion procedure from 6-week-old male Sprague-Dawley rats (Charles River, Saint-Germain-Nuelles, France) weighing 180–200 g, with the ethical committee authorization n°38096 (Ministère de l’Enseignement Supérieur et de la Recherche, France), as described previously [85]. Rat hepatocytes were seeded at 0.5×10^5^ cells/cm^2^ in William’s E medium supplemented with 10% Fetal calf serum, 100 units/mL penicillin, 100µg/mL streptomycin, 2mM L-glutamine, 5μg/mL insulin and 5.10^-7^M sodium hemisuccinate hydrocortisone. Adult hepatocytes were maintained either in pure culture or in coculture. Cocultures were prepared according to the conditions previously described [55, 86]. Briefly, 16 hours after hepatocyte seeding, the medium was discarded and RLEC SDVI cells and RLEC-like cell lines (S1, S2 and S3) established in this study (see section 2.7) suspended in fresh medium were added to hepatocyte cultures. At 24 hours, after RLEC cell adhesion, the medium was renewed and changed every 2 days thereafter.

### 4.2. Biopanning of PhD-12 Phage Display Library with HepaRG cholangiocyte cells

A biopanning of the Ph.D™.-12 M13 phage library expressing 12-amino acid sequences fused to the N-terminus of the minor coat protein PIII (New England BioLabs, Beverly, MA, USA) was performed using the Chol^DMSO^-HepaRG cells. This protocol was based on a whole cell subtractive approach. Negative/positive screen procedure was performed to obtain phages binding specifically to Chol^DMSO^-HepaRG cells. Hepatocyte-like HepaRG and HepG2 cells were used as negative absorber cells to eliminate phages interacting with hepatocytes and hepatoma cells (**I 1G**). Then, Chol^DMSO^-HepaRG cholangiocytes were used as final cell targets. During biopanning, the different cell types were washed 3 times with PBS and then blocked with serum-free medium containing 1% BSA for 1h. In each round of selection, ∼1.5×10^11^ pfu phages mixed to FCS free William’s E medium were incubated with hepatocyte-like HepaRG cells at 37°C for 1h under gentle agitation. The supernatant containing unbound phages was recovered and added to the blocked HepG2 cells under gentle agitation for 1h at 37°C. In the last step, the supernatant with unbound phages was collected, added to the blocked Chol^DMSO^-HepaRG cholangiocytes and incubated under gentle agitation for 1h at 37°C. The Chol^DMSO^-HepaRG cells were washed three times with PBS, once with PBS and Tween-20 at 0.1, 0.3, 0.5%, and a final wash with PBS to remove the unbound phages. The cell membrane-bound phages were then eluted with 800 µL of 0.2M glycine (pH 2.2) containing 1 mg.mL^-1^ BSA for 10 min on RT and immediately neutralized with 120μL of 1M Tris-HCl pH 9.1 (**SI 1G**). After each round of selection, the eluted phages were amplified and a small amount of the eluate was used for phage titration. For the amplification, the recovered phages were mixed with 25 mL of *E. coli* ER2738 strain culture at early log stage and incubated at 37°C with vigorous shaking for 4h. The culture was then centrifuged for 15min at 12,000g at 4°C. The upper 80% of the supernatant was pipetted to a fresh tube and mixed with 1/6 volume of 20% (w/v) PEG-8000 and 2.5M NaCl. The phages were precipitated overnight at 4°C. Following centrifugation at 10,000g for 10min at 4°C, the amplified phages were suspended in 200μL of TBS buffer solution (50mM Tris-HCl, 150mM NaCl, pH 7.5).

This procedure was repeated 3 times. After each round of selection, the eluted and amplified phages were mixed with the *E. coli* ER2738 strain and plated on LB medium agar plates containing isopropyl-beta-D-thiogalactoside (IPTG) (ICN Biomedical Inc., Brussels, Belgium) and 5-bromo-4-chloro-3-indolylbeta-D-galactopyranoside (Xgal) (Sigma-Aldrich, St. Louis MO, USA) for titration prior to the next round of selection. The binding efficiency of the phage pools were calculated by the ratio between output and input numbers in each round of screening. Phage plaques from the final round of panning were randomly picked out from tittered phage plaques for DNA extraction and sequencing (**SI 1G**).

### 4.3. DNA Sequencing of the Positive Phage Clones

The selected phage clones were used to extract DNA for sequencing analysis. An overnight culture of *Escherichia coli* ER2738 was diluted to 1:100 in 10mL LB medium and single β-galactosidase positive bacterial colonies were added. The mixture was shaken at 37°C for 4.5h, the supernatant was harvested following centrifugation of the bacteria at 13,000 rpm for 10 min and phage DNA was amplified and extracted following manufacturer’s instructions. Briefly, 200 μL of PEG/NaCl (2.5M NaCl with 20% [w/v] PEG-8000) were added to 1mL of supernatant to precipitate the phages. The precipitate was suspended in iodide buffer (10^−2^M Tris-HCl, pH 8.0, 10^−3^M EDTA and 4M NaI), and followed by ethanol precipitation at room temperature for 10 min. The single-stranded DNA (ssDNA) was recovered and dissolved in 50μL TE buffer (10^−2^M Tris-HCl, pH 8.0, 10^−3^M EDTA). DNA sequencing of the selected phages was carried out by the GATC Biotech company (Mulhouse, France) using the primer recommended by the Ph.D™-12 M13 phage library manufacturer (**SI 14**) and peptide sequences were deduced.

### 4.4. Phage affinity assay

Phages expressing full length peptides (12 amino acids) were selected for phage ELISA. Chol^DMSO^-HepaRG cholangiocytes were plated at a density of 10^4^ cells per well in 96-well plates. Three days later, the cell monolayers were washed with PBS and fixed with 4% paraformaldehyde in PBS for 15 min at 4 °C. Wells were then washed twice (PBS) for 10 min and incubated with blocking buffer (1% BSA in PBS) for 1h at 37°C. The phage clones were added to the wells at 2 concentrations 10^10^ and 10^8^ pfu/well and incubated with gentle agitation at 37°C for 1h. Blocking buffer was used as the blank control. Unbound phages were discarded and wells were washed 3 times with 0.1% Tween-20 in PBS (PBST). Then, phages were detected using 100μl of PBST containing 1μg of HRP-anti-M13 mouse monoclonal antibody (sc-53004, Santa Cruz Biotechnology, Dallas TX USA) added to each well and incubated under gentle agitation at 37°C for 1h. After 2 washes with 0.1% PBST, 200μl of freshly made 3,3’,5,5’–Tetramethylbenzidine (TMB, Sigma, Saint Louis MO USA) prepared by dissolving 4.5mg of TMB in 20mL of 50mM sodium citrate, pH 4.0, then supplemented with 35mL of 30% H_2_O_2_, were added to each well and kept at room temperature in the dark for 10min. Absorbance was measured at 370 and 630nm with a microplate spectrophotometer.

### 4.5. Peptide synthesis, peptide-cell binding assays by flow cytometry and confocal microscopy

The candidate peptides identified from the selected phage clones were synthesized (Eurogentec S.A, Liège, Belgium). In order to mimic the structure of peptide displayed on the phage, a spacer sequence Gly-Gly-Gly-Ser was added to the C-terminus of the peptide to block the negative charge of the C-terminal carboxylate as recommended by the phage library manufacturer. Peptides were also modified by adding a biotin moiety at the C-terminal end in order to set up a binding assay using fluorescent streptavidin. Peptide stock solutions were prepared in PBS at 1mM and stored at -20°C.

The biotinylated candidate peptides 10, 11, 12, 18, 46, 50 and 51 were bound to fluorescent Dylight^TM^488 streptavidin (Thermo Fisher Scientific, Waltham, MA, USA) for 30 min at 4°C. Then, the biotinylated peptide-Dylight^TM^488 streptavidin complexes were diluted in William’s E medium containing FCS to obtain different concentrations in peptides and streptavidin: 0.2μM peptide/0.6μM streptavidin, 0.4μM peptide/1.2μM streptavidin and 0.8μM peptide/2.4μM streptavidin. The complexes were incubated with cholangiocytes and different other cell types for 24h. After incubation, culture media were discarded, the cell monolayers were washed once with PBS and detached with trypsin and resuspended in William’s E medium containing FCS. The fluorescence intensity of 10,000 cells was analyzed with a Becton Dickinson le LSRFortessa™ X-20 (cytometry core facility of the Federative Research Structure Biosit, Rennes, France). Dot plots of forward scatter (FSC: x axis) and side scatter (SSC: y axis) allowed to gate the viable cells prior to detection the fluorescence emitted by streptavidin, Dylight^TM^ 488 conjugated using the FITC channel. Two parameters were analyzed: the percentage of positive cells and the intensity of fluorescence (mean of fluorescence) reflecting the binding of fluorescent peptides onto the cells and/or their cell uptake. Streptavidin-biotin binding was used as negative control. Flow cytometry data were analyzed using DIVA software (Becton Dikinson). Peptide-streptavidin complexes were also visualized by confocal microscopy in HepaRG cells and RLEC stained with Hoechst to detect nuclear DNA using a Leica TCS SP8 equipment and images were analyzed with Leica LAS software (Fluorescence microscopy core facility MRiC of the Federative Research Structure Biosit, Rennes, France).

### 4.6. Isolation of rat liver epithelial cells-like cells from non-parenchymal cell cultures

During the two-step collagenase perfusion of adult rat livers (section *4.2*), non-parenchymal liver cells were also isolated and cultured in William’s E medium supplemented with 10% Fetal calf serum, 100 units/mL penicillin, 100µg/mL streptomycin, 2mM L-glutamine prior to cell sorting. Non-parenchymal cells were maintained in culture for 4 to 7 days, then incubated with the biotinylated peptide 10 as negative control and P11^Chol^ bound to fluorescent Dylight^TM^ 488 streptavidin (Thermo Fisher Scientific, Waltham, MA, USA) for 30 min at 4°C. Then, the biotinylated peptide-Dylight^TM^ 488 streptavidin (0.6μM peptide/1.8μM) complexes were added to William’s E culture medium and incubated with non-parenchymal hepatic cells for 4h. Then, cells were detached with trypsin, resuspended in William’s E culture medium containing 20% FCS and sorted using Becton Dickinson FACSAria III equipment (cytometry core facility of the Federative Research Structure Biosit, Rennes, France). Peptide P11^chol^ positive cells, gated on FITC channel versus size (forward scatter, FSC), were collected in William’s E culture medium containing 20% FCS, plated at 0.5×10^5^ cells/cm^2^ and expanded in William’s E medium supplemented with 10% FCS prior freezing to establish a cell working bank.

### 4.7. Providencia stuartii adhesion and invasion assays

The clinical isolate of *Providencia stuartii* was obtained from the department of microbiology from the University Hospital Pontchaillou (Rennes, France) and was characterized for antibiotic resistances. This strain was resistant to erythromycin, ampicillin, amoxicillin + clavulanic acid but sensitive to kanamycin, gentamycin, and chloramphenicol antibiotics (**SI 7**). The bacteria adhesion and invasion assays were adapted from a previous study [58]. Bacteria were expanded in LB culture medium to reach 0.5 optical density, spun down and resuspended in William’s E medium prior to the infection for 30min at 37°C of progenitor and differentiated HepaRG cells as well as Chol^I/T/B^-HepaRG at different multiplicities of infection (MOI) ranging from 5 to 100 (**SI 7**). Since the number of HepaRG cells varied between the 3 different cultures conditions, the total number of bacteria during infection was adjusted to the number of cells. For the adhesion assay, cells monolayers were extensively washed with PBS prior to the bacteria extraction. For the invasion assay, after the 30 min of incubation at 37°C, the HepaRG cells were incubated in medium containing 100µg/mL gentamycin (Panpharma, Luitré-Dompierre, France) for 2h in order to eliminate extracellular bacteria. At the end of antibiotic treatment, culture medium was discarded and the HepaRG cells were lysed to release internalized bacteria with 200µL of 0.2% Triton X-100 (Amersham Pharmacia Biotec, France) in PBS incubated for 10 min at room temperature. Subsequently, 20µl of serially diluted bacterial suspensions were plated on trypticase soy agar, incubated for 24h at 37°C, and bacteria colonies were counted. The results were presented as the number of colonies per 10^5^ HepaRG cells. For experiments with peptides, cells were incubated with peptides 10 and P11^Chol^ at 10 and 25µM culture medium at 37°C for 1h prior to infection with *P. stuartii* suspension with a cell/bacteria ratio of 1:10 at 37°C for 30min.

A recombinant *Providencia stuartii* expressing GFP was also generated by transformation with the modified plasmid pNF8 (pNF8 *gfp-mut1*) [87] to visualize GFP positive bacteria during adhesion and internalization assay with HepaRG cells by confocal fluorescence microscopy. The laser outputs are controlled via the Acousto-Optical Tunable Filter (AOTF) and photomultiplicators (PMT) as follows: Hoechst 33342 was excited with a laser diode at 405 nm (17-22%) and measured with an emission setting at 410-443 nm; GFP was excited with an Argon laser at 488 nm (15-20%) while the emission was collected in the 495-544 nm range. Transmission images were collected on the transmission photomultiplicators (T-PMT) using Helium-Neon laser at 633 nm in transmission mode.

### 4.8. Cell viability analysis by caspase assay and ATP quantification

Caspases activities were quantified as previously described [88]. Briefly, cells were lysed in caspase-activity buffer containing PIPES (20mM: pH 7.2), NaCl (100mM), EDTA (1mM), CHAPS (0.1%), sucrose (10%) and homogenized by sonication. Fifty µg of total proteins were added to caspase-activity buffer supplemented with 10mM DTT and 80μM of substrate Ac-DEVD-AMC (ALX 260-031) purchased from Enzo Life Science (Farmingdale, NY, USA) to measure caspase 3 activities. The luminescence was measured using the Polarstar Omega microplate reader (BMG Labtech, Champigny sur Marne, France, excitation/emission wavelengths at 380 and 440 nm) during a 2h-time course with end-point representation data. Cell viability was evaluated in progenitor and differentiated HepaRG cell cultures after *Providencia stuartii* adhesion/internalization assays by measuring the intracellular ATP content using the CellTiter-Glo Luminescent Cell Viability Assay (Promega, Madison, WI, USA) according to the manufacturer’s instructions. Briefly, untreated and treated HepaRG cells were first lysed with the CellTiter-Glo reagent for 10 min at 37°C. Cell lysates were then transferred in opaque-walled 96-well plates and the luminescent signal was quantified at 540 nm with the POLARstar Omega microplate reader (BMG Labtech, Champigny sur Marne, France). ATP levels in treated cells were expressed as the percentage of the ATP content measured in untreated cells.

### 4.9. Real-time quantitative Reverse Transcription PCR (RT-qPCR) analysis

Total RNAs were purified from HepaRG cells using a Macherey-Nagel NucleoSpin^®^ RNAII kit, according to the manufacturer’s protocol. Following elution, RNA quantity and purity were assessed with a Nanodrop ND-1000 spectrophotometer (Nyxor Biotech, Paris, France). Total RNAs (0.5μg) were reverse-transcribed into first-strand cDNA using a High-Capacity cDNA Achieve Kit (Applied Biosystems, Foster City, CA, USA), according to the manufacturer’s instructions. Real-time qPCR was performed using the fluorescent dye SYBR Green method, with SYBR Green PCR Master Mix in 384-well plates and the StepOnePlus^TM^ system (Applied Biosystems, Thermo Fisher Scientific, Waltham, MA, USA) using the ABI PRISM 7900HT instrument. TATA binding protein was used as the reference gene. Relative quantification values were expressed using the 2^ΔΔCt^ method as fold changes in the target gene, normalized to the TATA binding protein reference gene, and related to the expression level in control experiments (arbitrarily set to 1). The sequences of the forward and reverse primers are provided in **SI 14.**

### 4.10. Phase I and II enzymatic activities and bile salt transport

RLEC lines, hepatocyte pure cultures and cocultures with RLEC were incubated for 1h with a mix of phase I probe substrates, i.e. bupropion (100µM) purchased from Sigma-Aldrich, and midazolam (100µM) obtained from Pharmacopée Européenne, to quantify CYP2B6 and CYP3A4/5, respectively. CYP2D6 activities were evaluated by CL_int_ method using dextromethorphan as substrate purchased from Sigma-Aldrich). Phase II enzymatic activities (sulfotransferases and uridine 5′-diphospho-glucuronosyltransferases) were investigated by measuring the biotransformation of 1mM paracetamol via the production of sulfate- and glucuronide-paracetamol conjugates. The reactions were stopped by addition of ice-cold acetonitrile and samples were analyzed (10µL) by liquid chromatography coupled to tandem Mass Spectrometry (LC/MS-MS). The analysis was performed on an Exion LC AC system (Applied Biosystems/Sciex, USA). Compounds were separated on a Xselect HSS T3 column (2.1×75mm, 2.5µm, Waters, Milford, MA) maintained at 35°C. The mobile phase was composed of 2 mM ammonium formate containing 0.1% formic acid (A) and acetonitrile (B). Detection was operated on a triple quadrupole mass spectrometer, AB Sciex Triple Quad 5500+ (Applied Biosystems/Sciex, USA), equipped with an electrospray ion source in positive mode. Acquisition was performed using the multiple reaction monitoring (MRM). Data processing was performed using Analyst 1.7.2 software.

The uptake and clearance of [^3^H]-taurocholic acid (6.7 Ci/mmol, 37 MBq/mL, PerkinElmer Waltham, MA USA) was studied in confluent RLEC SDVI and RLEC-like S1 and S2 cell lines, 21 days after plating. Briefly, RLEC cells were incubated for 30 min with 75 kBq of [^3^H]-taurocholic acid corresponding to a final concentration of 40nM in bile salt to evidence bile salt accumulation. After discarding medium containing [^3^H]-taurocholic acid, RLEC cultures were washed 3 times with PBS before cell lysis with NaOH 0.1M. In other culture wells, fresh medium was added after the 30min incubation with [^3^H]-taurocholic acid and clearance of the bile salt was studied at 1 and 4h. All cell lysates were transferred into vials and counted using A TriCarb liquid scintillation analyser (PerkinElmer Waltham, MA USA). Results were expressed as disintegrations/min (CPM) per well.

The active transport of bile salts by functional bile canaliculi of hepatocytes in cocultures and pure cultures was visualized at day 7 after plating using cholyl-lysyl-fluorescein (CLF, reference 451041, Becton Dikinson, Franklin Lakes NJ, USA), which accumulates inside hepatocyte neocanaliculi [60]. CLF was added in culture medium at 5μM CLF and incubate 15 min at 37°C, then medium was discarded and coculture were washed twice. Fresh phenol red free medium was added and fluorescence was detected using a Zeiss Inverted Microscope and Axio-Vision Software (Carl Zeiss AG, Oberkochenn Baden-Württemberg, Germany).

### 4.11. Microbiota 16S analysis in healthy and tumoral human livers

Total DNA was extracted from 13 healthy human liver biopsies from patients subjected to surgical resections for hepatic colorectal metastasis and 17 paired biopsies of both non-tumoral liver parenchyma and CCK. Patients (males and females) were from 51 to 78 years old with means at 62.2 ± 8.4 and 63 ± 7.1 for healthy and CCK patients, respectively (**SI 15).** DNA was extracted from frozen biopsies using the DNeasy blood and tissue kit (Qiagen, Hilden, Germany). The V3-V4 region of 16S rRNA gene was amplified using specific primer sequences (**SI 14**) and sequenced using Illumina Miseq technology. The raw sequences were analyzed using QIIME 2 bioinformatic pipeline (v. 2018.4, https://qiime2.org/) as previously described [87]. The analysis of Core diversity was performed with QIIME2 diversity core-metrics-phylogenetic plugin, with specific 11399 sampling depth.

Total RNAs were also purified from the same human liver biopsies using a Macherey-Nagel NucleoSpin^®^ RNAII kit and were reverse-transcribed into first-strand cDNA as described in section 2.10. The quantification of bacterial 16S RNA expressed in *Providencia*, *Photorhabdus* and *Salmonella enterica* species among the total liver RNAs was performed by RT-qPCR as described in section 2.10 using reverse-transcribed cDNAs and forward/reverse 16S primers (PsPrSe-16S) designed from conserved RNA sequences for these bacteria species (**SI 14**). Expression of bacterial 16S RNA was normalized to the human Hypoxanthine Phosphoribosyltransferase 1 (HPRT) reference gene.

### 4.12. Protein alignments and structure predictions

Protein homology analysis and sequence alignment were performed using Blast protein Swissprot all genomes and Cobalt open access websites at the National Center for Biotechnology Information NCBI (National Institute of Healthy-NIH, Rockville Pike, Bethesda, MA USA) and refined manually. The crystal structures of BamA/TamA from *Salmonella enterica* subspecies serovar Typhimurium and *Escherichia coli* (*E. coli*) were available from NCBI’s Molecular Modeling Database (MMDB) [89]. TamA structure references from *Salmonella enterica* are PDB ID 5OR1 and MMDB ID 159100 [90]. TamA structure references from *E. coli* are: PDB ID 4C00 and MMDB ID 113694 [63]. Structure of BamA from *Providencia stuartii* was predicted using iCn3D version 4.6 macromolecular structure viewer (NCBI).

### 4.13. Statistical analysis

The results were expressed as mean ± standard error of the mean (SEM) of 2 to 6 independent experiments (n= 2 to 6) including 3 to 6 independent culture wells in each experiment. Comparisons between groups were performed using one-way ANOVA followed by a post hoc Dunnett’s test or Bonferroni’s test when the normality test was positive, and Kruskal-Wallis followed by a post hoc Dunn’s test when data were not normally distributed. Graph Pad Prism 8 software (GraphPad Software, San Diego CA, USA) was used for all statistical analyses and the threshold for statistical significance was *\*p value* < 0.05, \*\**p* < 0.01 and \*\*\**p* < 0.001. In experiments of *Providencia stuartii* adhesion and invasion assays of human HepaRG cholangiocytes, paired samples *t*-tests (dependent samples *t*-tests) were used. For bacterial 16S RNA analysis, species richness (Chao1) and evenness (Shannon index) were calculated for alpha diversity estimations and compared using Mann-Whitney tests, between each group. A *p* value < 0.05 was considered significant. Taxonomy composition comparison were performed with Mann-Whitney test between each group.

## 5. Acknowledgements

The authors would like to thank Dr. Agnès Burel from the Mric microscopy core facility (Federative Research Structure Biosit, Rennes, France) for helping with transmission electron microscopy. The authors would also like to thank Dr. Nicolas Crouvezier, project manager of the valorization organization Inserm transfer for his assistance in filing the invention declaration and writing the patent describing Peptide P11^Chol^ and its applications.

## 6. Statements and declarations

### Ethics statement and clinical trial

Healthy human liver biopsies, cholangiocarcinomas (CCK), primary human hepatocytes (PHH) and non-parenchymal liver cells were obtained from the processing of biological samples through the Centre de Ressources Biologiques of Rennes (CRB Santé of Rennes, France, Institutional review board approval BB-0033-00056) and the Réseau National des CRB Foie (BB-033-00085) under French legal guidelines and fulfilled the requirements of the institutional ethics committee including informed consent statement collected by the Centre de Ressources Biologiques (CRB Santé of Rennes, France). Clinical trial number: not applicable.

### Data Availability Statement

Data supporting reported results will be publicly available at Mendeley Data [https://data.mendeley.com/] and accessible using the following link: LOYER, Pascal (2024), “A novel peptide derived from Enterobacteriaceae BamA/tamA-like outer membrane proteins inhibits Providencia stuartii internalization by cholangiocytes in vitro and provides a new mean to isolate liver primary epithelial cells.”, Mendeley Data, V1, doi:10.17632/nxrybyg3kh.1.

### Patent

The peptides P11^Chol^ [seq ID: TFLNSVPTYSYWGGGS], Provid-Stu [seq ID: AFLNSNGLPTYSPKIS] and Hel-Pyl [seq ID: GTLNNIPTYYWGKGY] and derived peptides were patented by Inserm Transfert (PariSanté Campus, 75015 Paris) as a European Patent on August 26, 2025, under the registration number EP25306338.

### Author’s Contribution

Conceptualization, P.M., L.B., D.O., A.C. and P.Lo.; methodology, S.D-L.G., L.B., A.C. and P.Lo.; validation, S.D-L.G., L.B., A.C. and P.Lo.; formal analysis, S.D-L.G., A.C. and P.Lo.; investigation, P.M., C.R., P.Le, M.V., H.C-E., K.M., H.N., N.S. and P.Lo ; data curation, P.M. and P.Lo.; writing—original draft preparation, P.M. and P.Lo.; writing—review and editing, P.M., S.D-L.G., L.B., A.C. and P.Lo. ; supervision, D.O., A.C. and P.Lo.; project administration, P.Lo.; funding acquisition, P.M., H.C-E., K.M. and P.Lo. All authors reviewed the submitted version of the manuscript.

## 9. Supporting Information Captions

**Supporting Information 1. HepaRG cell model and the biopanning of Chol^DMSO^-HepaRG cells**. Progenitor cells actively proliferate for ∼10 days after seeding at low density (A-B) and produce a confluent monolayer in ∼14 days (C). At this stage, the cells have committed to differentiation towards hepatocyte (Hep) or cholangiocyte (Chol) pathways. The differentiation is further enhanced by adding DMSO at 2% to the culture medium over a period of 2 weeks resulting in a coculture of both cholangiocytes (Chol^DMSO^-HepaRG cells) and hepatocytes (D). The Chol^DMSO^-HepaRG cells (F) were isolated by selective detachment of hepatocyte-like HepaRG cells, which were also used to produce a pure culture of hepatocytes (E) and to perform 2 subtractive biopannings of the phage library on a non-cholangiocytic hepatic target (G). The biopanning also included subtractive binding of the remaining phages using HepG2 cells prior to the final biopanning with the targeted Chol^DMSO^-HepaRG cells. Three rounds of biopanning were performed with amplification and titration of the eluted phages from the Chol^DMSO^-HepaRG cells after each panning. At the third round, the DNA of 50 individual clones of phages was sequenced to identify the random sequence inserted into the protein PIII of the M13 phage.

**Supporting Information 2. Results of biopanning and identification of peptide sequences.** A) Input versus output of Pfu ratio obtained during the biopanning. For each round of biopanning, 1.5×10^11^ Pfu were incubated in the first well of hepatocyte-like HepaRG cells used for the subtractive panning (S 1G), then unbound phages were transferred to a second well of hepatocyte-like HepaRG cells and 2 wells of HepG2 cells prior binding to Chol^DMSO^-HepaRG cells. The output corresponds to the numbers of phages eluted from Chol^DMSO^-HepaRG cells determined by titration The titers of eluted phages increased between the first and the third rounds indicating the enrichment in phages of higher affinity for Chol^DMSO^-HepaRG cells. B) List of phage clones isolated during the selection procedure, the amino acid sequences of full-length peptides (12 amino acids) obtained by phage genome sequencing and translation, and frequency of occurrence of clones. C) Elisa phage affinity assay on the Chol^DMSO^-HepaRG cell target. The 7 peptides 10, 11, 12, 18, 46, 50 and 51 were selected and synthesized for affinity tests. The background baseline was set at an optical density (OD) value of 1.5 in two independent experiments (n=2).

**Supporting Information 3. Synthetic biotinylated peptides used for cell binding assay.** A) List of selected peptides, number of copies of corresponding phages found in the screen, peptide selection parameters (ELISA affinity and/or phage frequency and peptide sequences synthesized for affinity assays) and synthetic peptide sequences. At the c-terminus end of the 12 amino acid peptides, a spacer sequence Gly-Gly-Gly-Ser (GGGS) was added to block the negative charge of the C-terminal carboxylate as recommended by the phage library manufacturer. Peptides were also modified by adding a biotin moiety at the C-terminal end in order to set up a binding assay using fluorescent streptavidin. B) Scheme of the cell-peptide interaction assay based on the use of synthetic biotinylated peptides and bound to fluorescent Dylight^TM^488 streptavidin (SA) before incubation of the complex with the cells in culture and fluorescence analysis by flow cytometry. Negative control corresponded to fluorescent streptavidin saturated with biotin.

**Supporting Information 4. Binding of peptide P11^Chol^ to different cell types.** Fluorescence histograms obtained for the binding of peptide 11 with different hepatic cell types (HepG2, LX2, HepaRG-SP, HuGB, RLEC, Chol^DMSO^- and Chol^I/T/B^ HepaRG cells) or non-hepatic cells (NCI-H727, HEK293T). The dot plots (FSC-A) versus structure (SSC-A) on the left show the viable cell populations on which the gates were set to measure the fluorescence (FITC-A) of the fluorescent peptide 11-Dylight^TM^488 streptavidin complex represented on the abscissa axis of the right histograms according to the analyzed cell number (counts, y axis).

**Supporting Information 5. Cell internalization of peptide P11^Chol^-streptavidin complex by endocytosis.** A) Confocal microscopy of the internalized peptide 11-Dylight^TM^ 488 streptavidin complexes in Rat Liver Epithelial Cells (RLEC), Chol^DMSO^-HepaRG cells. Biotinylated peptide was coupled to fluorescent Dylight^TM^ 488 streptavidin (green) and the nuclear DNA was detected with Hoechst (blue). The peptide 10-Dylight^TM^488 streptavidin complex was used as control for background fluorescence. B) Flow cytometry histogram overlay and C) quantification of fluorescence intensities (FITC-A mean) measured in Chol^DMSO^-HepaRG cells incubated with biotin-Dylight^TM^ 488 streptavidin complex (background fluorescence) or peptide 11-Dylight^TM^ 488 streptavidin complex in absence or in presence of endocytosis inhibitors genistein (200 mM) and chlorpromazine (20 mM).

**Supporting Information 6. The main physical and chemical characteristics of the peptide 11.** Using ProtParam tool for primary structure analysis from ExPASy bioinformatics database (https://web.expasy.org), the peptide 11 molar weight (Mw) is 1735.87 g/L with an isoelectric point (pI) at 5.18 while peptide 10 exhibits a theorical Mw of 1819.02 g/L and a pI of 6.07. Peptide 11 contains a relatively low percentage on hydrophobic amino acids with GRAVY value at -0.175 indicating a low hydrophobicity in agreement with its high solubility in aqueous solution. The instability index was 31.59 that classified the peptide as stable and the estimated half-life was 7.2h in mammalian reticulocytes.

**Supplementary Information 7. *Providencia stuartii* adhesion and internalization assay in HepaRG cells. A)** Clinical isolate of *Providencia stuartii* was used to adapt an adhesion and internalization assay in Chol^I/T/B^-HepaRG cells. The *Providencia stuartii* strain was resistant to erythromycin, ampicillin, amoxicillin+clavulanic acid but sensitive to kanamycin, gentamycin, and chloramphenicol antibiotics. Bacteria were expanded in LB culture medium to reach 0.5 optical density, spun down and resuspended in William’s E medium prior to the incubation for 30 min at 37°C with progenitor and differentiated HepaRG cells as well as Chol^I/T/B^-HepaRG at different multiplicity of infection (MOI) ranging from 5 to 100, with bacteria numbers adjusted to the number of cells. **B**) For the adhesion assay, cells monolayers were extensively washed with PBS prior to the bacteria extraction. For the invasion assay, after the 30 min of incubation at 37°C, the HepaRG cells were incubated in medium containing gentamycin (100µg/ml) for 2h in order to eliminate extracellular bacteria. At the end of antibiotic treatment, culture medium was discarded and the HepaRG cells were lysed to release internalized bacteria with 200µl of 0.2% Triton X-100 (Amersham Pharmacia Biotec, France) in PBS incubated for 10 min at room temperature. Subsequently, 20µl of serially diluted bacterial suspensions were plated on trypticase soy agar, incubated for 24h at 37°C, and bacteria colonies were counted. The results were presented as the number of colonies per 10^5^ HepaRG cells. **C**) Recombinant strain of *Providencia stuartii* expressing the Green Fluorescent Protein (GFP) was generated to visualize bacteria by confocal microscopy after the adhesion and internalization assay in Chol^I/T/B^-HepaRG cells, which were stained with Hoechst to visualize the DNA. GFP^+^ bacteria bound to the cell (adherent bacteria) were detected out of the focal plane of nuclei (white arrows) and internalized bacteria (green arrows) were found on the same focal plane than cell’s nuclei.

**Supplementary Information 8. A)** Impact of the infection by *Providencia stuartii* at different MOI on the cell viability in cultures of progenitor and DMSO-differentiated HepaRG cells as wells as Chol^I/T/B^-HepaRG cells by measuring the relative intracellular content in ATP after the incubation with the bacteria. B) Ultrastructure of control Chol^I/T/B^-HepaRG cells and cells incubated with *Providencia stuartii* for 30 min, washed with PBS, prior to fixation with glutaraldehyde and further process for TEM. Dark arrows delineate nuclei (N), plasma membrane (PM), bacteria (Bact), autophagosomes (Autoph) and large lysosomes (Lyso).

**Supplementary Information 9.** Flow cytometry analysis of binding assays using biotin control- and peptide-Dylight^TM^488 streptavidin complexes and rat liver epithelial cells (RLEC SDVI). Cells were incubated with negative biotin control-Dylight^TM^488 streptavidin complex (A), and peptides 10 (B), P11^Chol^ (C), Provid-Stu (D) and HelPyl (E) bound to Dylight^TM^488 streptavidin as described in Supplementary Information 6. Cells were analyzed by FACS to gate viable cells using SSC-A versus FSC-A (dot plots on left panels and gate P1). Then, single cells were visualized using the SSC-A versus SSC-H (second dot plots and P2 gate). FITC fluorescence was analyzed on single cells from P2 gate and plotted as histograms (cell count versus FITC) and dot plots (SSC-A versus FITC). Gates P3 and P7: negative cells, gates P4 and P8: positive cells.

**Supplementary Information 10.** Photographs in phase contrast microscopy illustrating the morphologies of RLEC SDVI and RLEC-like lines isolated in this study at different time points after seeding. All cells were plated in regular plastic dishes at density of ∼5.10^5^ cells/cm^2^. Cells rapidly attached after plating to form small RLEC colonies at day 1. They proliferate actively during the first week after seeding with an estimated doubling time of ∼30 hours. At day 7, all RLEC lines form a confluent monolayer of quiescent cells. At day 14, colonies with heterogenous morphologies can be observed suggesting a process of differentiation. When maintained confluent over several weeks, RLEC remain viable and stable with production of an abundant extracellular matrix visible in phase contrast microscopy (high magnification at day 60). At this stage, while most of the cell populations form two-dimensional monolayers, spherical cell/cysts structures can be observed suggesting formation of polarized three-dimensional architecture for a limited number of cells even when RLEC are plated on plastic support.

**Supplementary Information 11.** Detection of CD133, CD24 and Cystic Fbrosis Transmembrane Conductance Regulator (CFTR) by flow cytometry in RLEC SDVI and RLEC-like cell lines established in this study. A) Isotype and anti-CD133 histograms in RLEC SDVI and RLEC-like cells: CD133 was not detected in RLEC-like S2 and S3 cells while anti-CD133 fluorescence signal was slightly higher in RLEC SDVI and RLEC-like S1 cells compared to mean of fluorescence obtained with isotype antibodies suggesting a low CD133 expression in RLEC SDVI and RLEC-like S1 cells. In contrast, all RLEC lines were strongly positive for CD24 (B) and CFTR (C) expression.

**Supplementary Information 12.** Photographs in phase contrast microscopy illustrating the morphology of hepatocytes in pure culture at 24 (A), 72 (B) hours (h) and after 7 days of culture, and in coculture with rat liver epithelial cells (RLEC) upon addition RLEC cells (D), 72 h (E) and 7 days (F) after addition of RLEC cells. Scale bar: 100 mm. After plating, hepatocyte rapidly attach to tissue culture dishes to from well-defined colonies. hepatocytes characterized by a cuboidal shape, a dark cytoplasm, a large nucleus with a single nucleolus (A). In pure culture, hepatocytes progressively spread on plastic dishes and lose their typical morphology (B, C) : the cytoplasm becomes clearer, the nuclei enlarge and the nucleoli are fragmented. After days 3 to 5, some cells detach characterizing progressive cell death by apoptosis [77]. In coculture, RLEC are seeded within 24 h after hepatocyte plating (D), which attach on plastic dish and spread between hepatocyte colonies. Hepatocytes keep their typical morphology with dark cytoplasm, well-defined nuclei (N) and plasma membrane (PM) and bile neo-canaliculi (BCN).

**Supporting Information 13.** Analysis of functional bile canaliculi in hepatocytes using cholyl-lysyl-fluorescein (CLFI) efflux, detection in fluorescence microcopy and quantification of the canaliculus network with ImageJ software and the free macro *Tool Scale Bar Tools for Microscope* (http://image.bio.methods.free.fr/ImageJ/?Scale-Bar-Tools-for-Microscopes.html) designed by Gilles Carpentier (Faculté des Sciences et Technologie, Université Paris Est, Créteil Val-de-Marne, France). Length of bile canaliculi (white bars and arrows) were measured for colonies including at least 6 hepatocytes or more. Hepatocytes were counted in each colony and ratio of canaliculi/number of hepatocytes was established. Results were obtained from 4 independent experiments in which at least 6 images (fields of cocultures, with a total of 250 to 500 hepatocytes) were analyzed.

**Supporting Information 14.** Primer sequences

**Supporting Information 15.** Gender and age of healthy versus CCK patients from whom liver biopsies were used in the study.

