## Supporting information for "Isolation of rat and human hepatic cholangiocytes using a peptide derived from a conserved domain of enterobacteria BamA/TamA-like proteins"

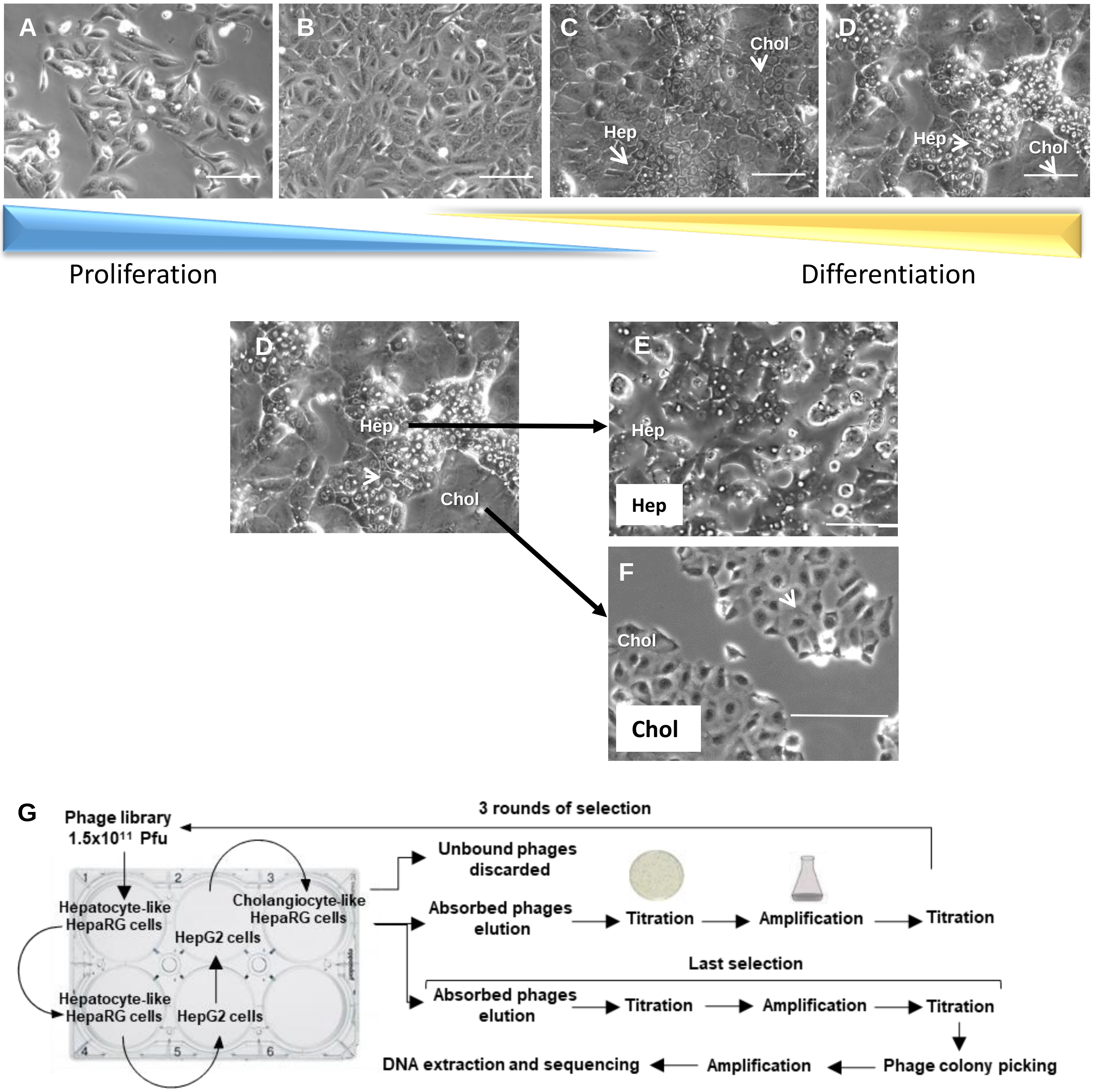


**Supporting Information 1**

**A**

| Input (Pfu) | |
|---|---|
| **Round 1** | $1.5 \times 10^{11}$ |
| **Round 2** | $1.35 \times 10^{11}$ |
| **Round 3** | $1.5 \times 10^{11}$ |
| **Output (Pfu)** | |
| **Round 1** | $6.48 \times 10^{4}$ |
| **Round 2** | ND |
| **Round 3** | $9.54 \times 10^{5}$ |
| **Output/Input** | |
| **Round 1** | $4.32 \times 10^{-7}$ |
| **Round 2** | ND |
| **Round 3** | $6.35 \times 10^{-6}$ |

**B**

| Phage Number | Amino acid sequence (PhD™-12 library, New England BioLabs® Inc.) | Frequency |
|---|---|---|
| Clone 1 | TIVENHYQTHIL | 1/33 |
| Clone 5 | YAFTSEISYTMF | 1/33 |
| Clone 7 | INSTPNPKMFPS | 1/33 |
| Clone 8 | RSTLPVDHLIRS | 1/33 |
| Clone 10 | YTTKEFLMRWDA | 2/33 |
| Clone 11 | TFLNSVPTYSYW | 1/33 |
| Clone 12 | GTELRDGEHRSH | 2/33 |
| Clone 18 | TTDFDKHSRFPS | 14/33 |
| Clone 31 | TDTHIVTYMSLA | 1/33 |
| Clone 32 | YLDVVGLTTMLH | 1/33 |
| Clone 35 | GTDIIHPRVIFN | 1/33 |
| Clone 38 | TIPTRDPAMLHS | 1/33 |
| Clone 46 | GAISTYSNRFLP | 1/33 |
| Clone 47 | SEHLLAMYTSYS | 1/33 |
| Clone 50 | AGHSVWSIEKQY | 2/33 |
| Clone 51 | YSASTLSTKLSE | 2/33 |

**C**

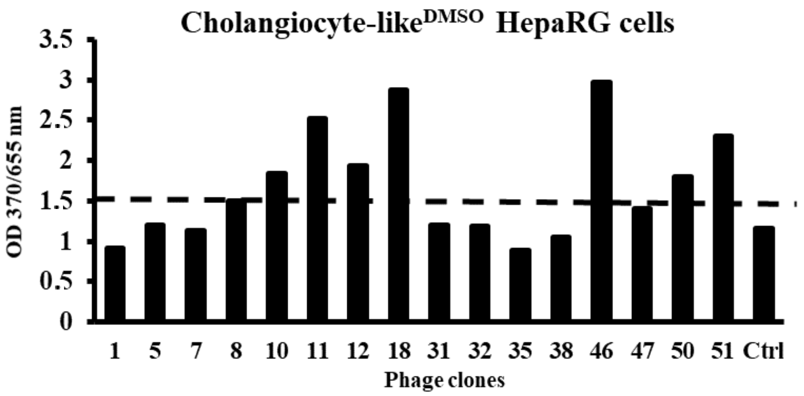


**Supporting Information 2**

**A**

| Peptide Number | Copy Number (panning) | Selection : Copy number/Elisa | Amino acid sequences |
|---|---|---|---|
| 10 | 2 | Elisa/copy number | Y T T K E F L M R W D A G G G S |
| 11 | 1 | Elisa | T F L N S V P T Y S Y W G G G S |
| 12 | 1 | Elisa | G T E L R D G E H R S H G G G S |
| 18 | 14 | Elisa/copy number | T T D F D K H S R F P S G G G S |
| 46 | 1 | Elisa | G A I S T Y S N R F L P G G G S |
| 50 | 2 | Elisa/copy number | A G H S V W S I E K Q Y G G G S |
| 51 | 2 | Elisa/copy number | Y S A S T L S T K L S E G G G S |

**B**

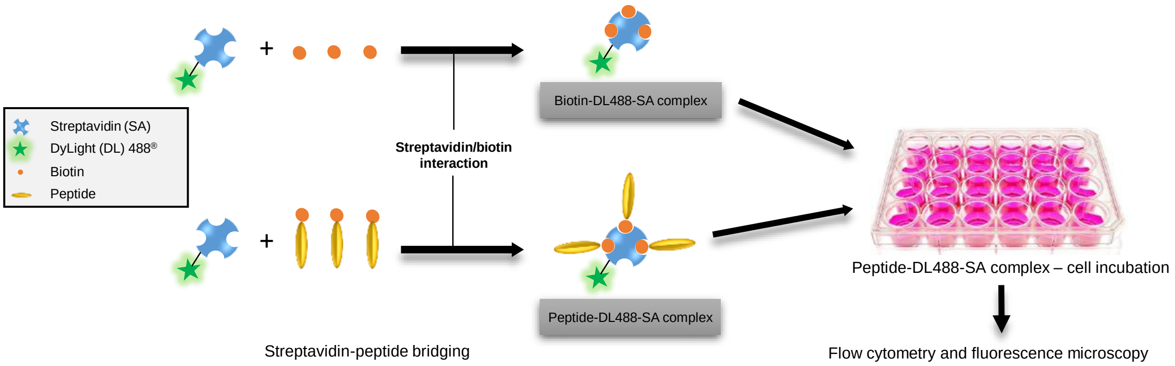


**Supporting Information 3**

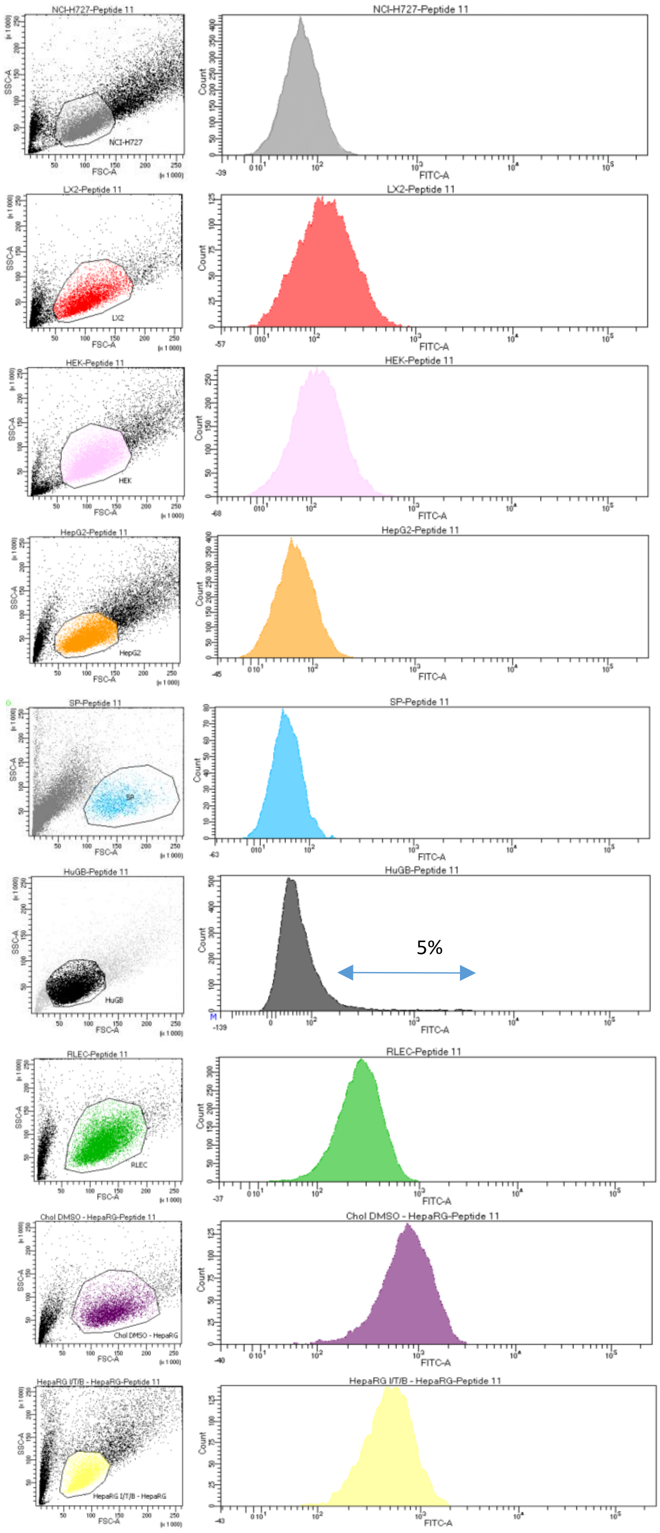


**Supporting Information 4**

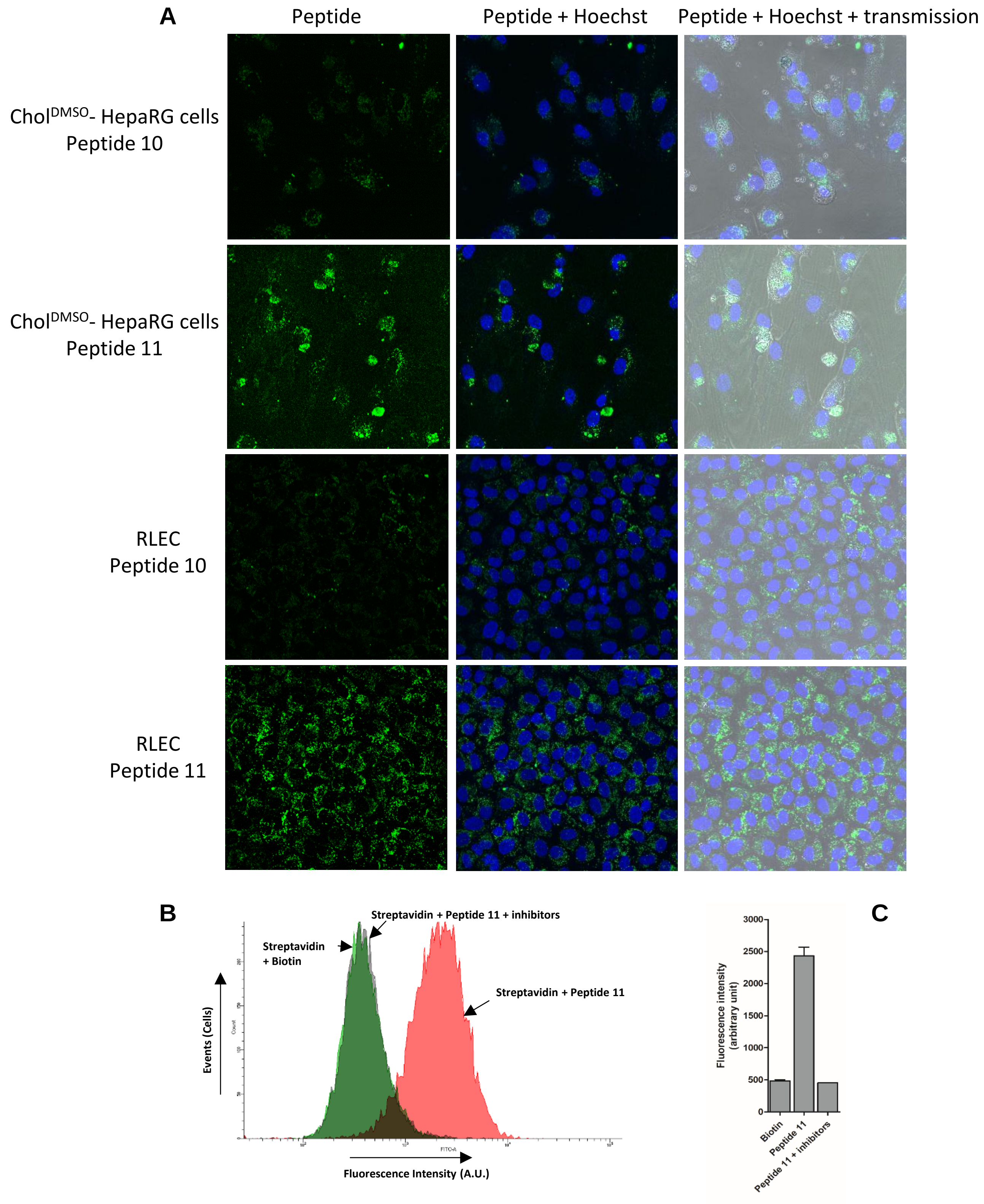


**Supporting Information 5**

| Peptides | Theoretical Mw | Theoretical pI | Ext. coefficient | Estimated half-life (hours) | Instability index | Aliphatic index: | Hydropathic (GRAVY) |
|---|---|---|---|---|---|---|---|
| 10 | 1819.02 | 6.07 | 6990 | 2.8 | 50.58 | 30.63 | -0.669 |
| 11 | 1735.87 | 5.18 | 8480 | 7.2 hours | 31.59 | 42.50 | -0.175 |

Peptide 11 composition :

```
Asn (N)   1     6.2%
Gly (G)   3    18.8%
Leu (L)   1     6.2%
Phe (F)   1     6.2%
Pro (P)   1     6.2%
Ser (S)   3     18.8%
Thr (T)   2     12.5%
Trp (W)   1     6.2%
Tyr (Y)   2     12.5%
Val (V)   1     6.2%
```

**Extinction coefficients:**
Extinction coefficients are in units of $M^{-1}$ $cm^{-1}$, at 280 nm measured in water.
Ext. coefficient 8480
Abs 0.1% (=1 g/l) 4.885

**Estimated half-life:**
The estimated half-life is: 7.2 hours (mammalian reticulocytes, *in vitro*), 20 hours (yeast, *in vivo*), 10 hours (*Escherichia coli*, *in vivo*).

**Instability index:**
The instability index (II) is computed to be 31.59
This classifies the peptide as stable.

**Aliphatic index:** 42.50

**Grand average of hydropathicity (GRAVY):** -0.175

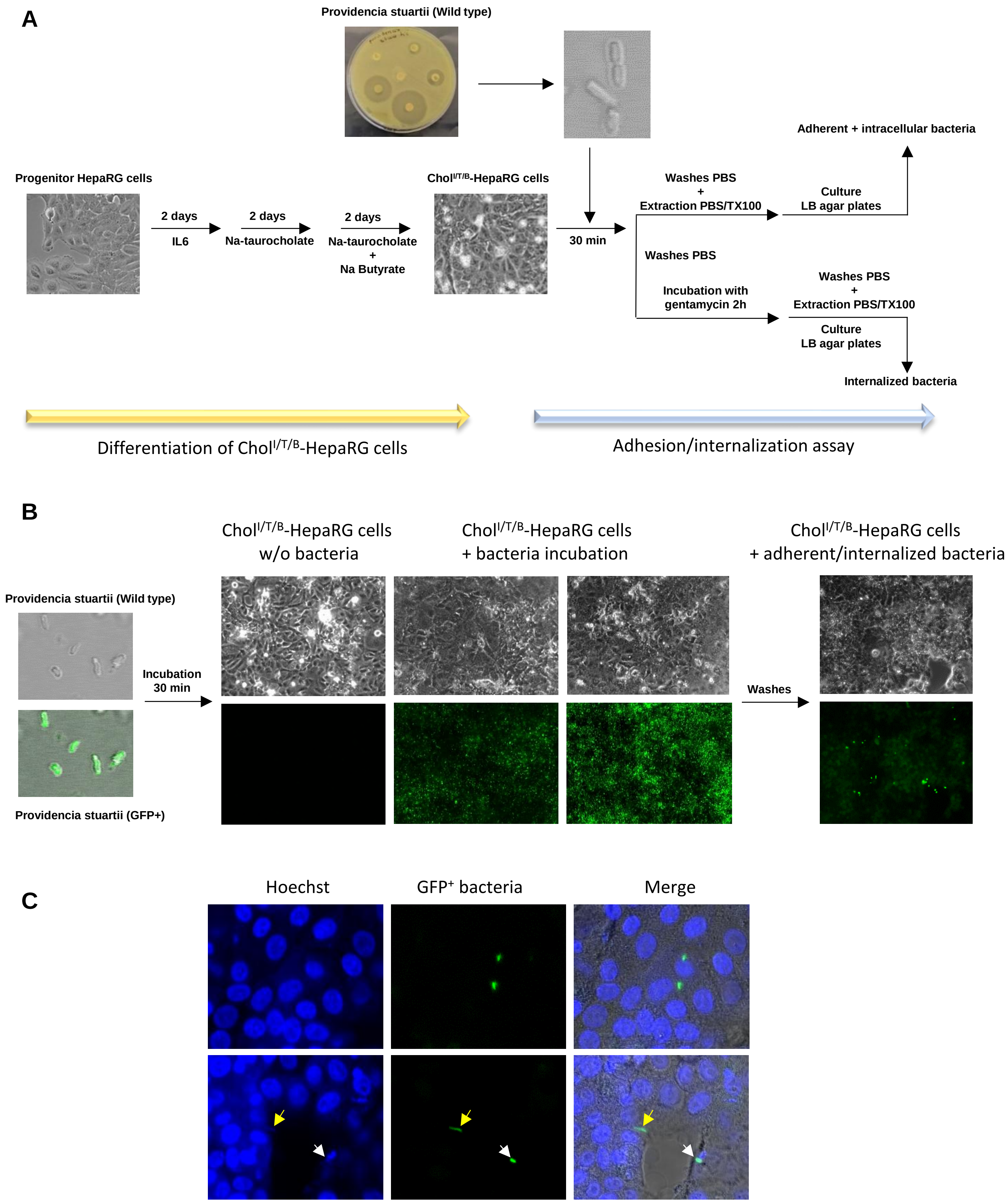


**Supporting Information 7**

**A**

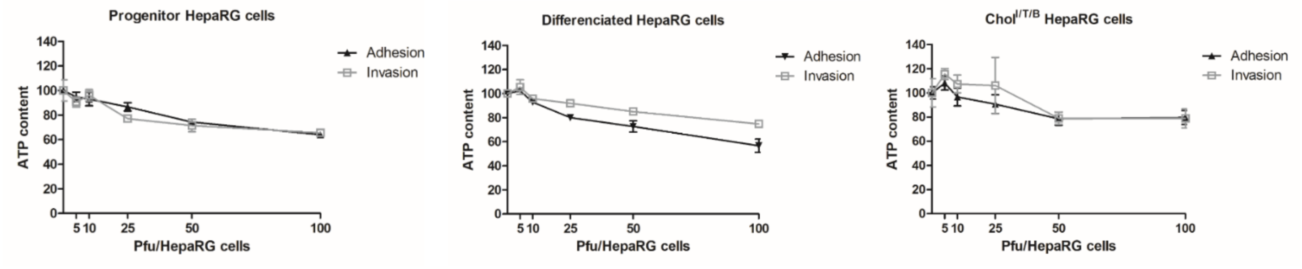


**B**

Control chol$^{I/T/B}$-HepaRG cells w/o bacteria

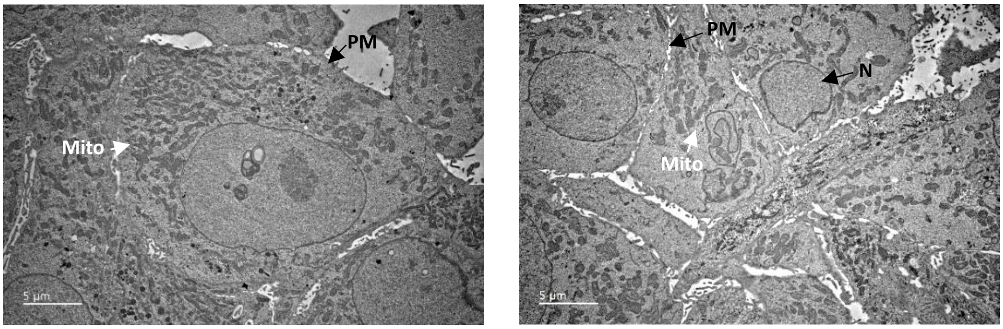


Chol$^{I/T/B}$-HepaRG cells incubated with bacteria

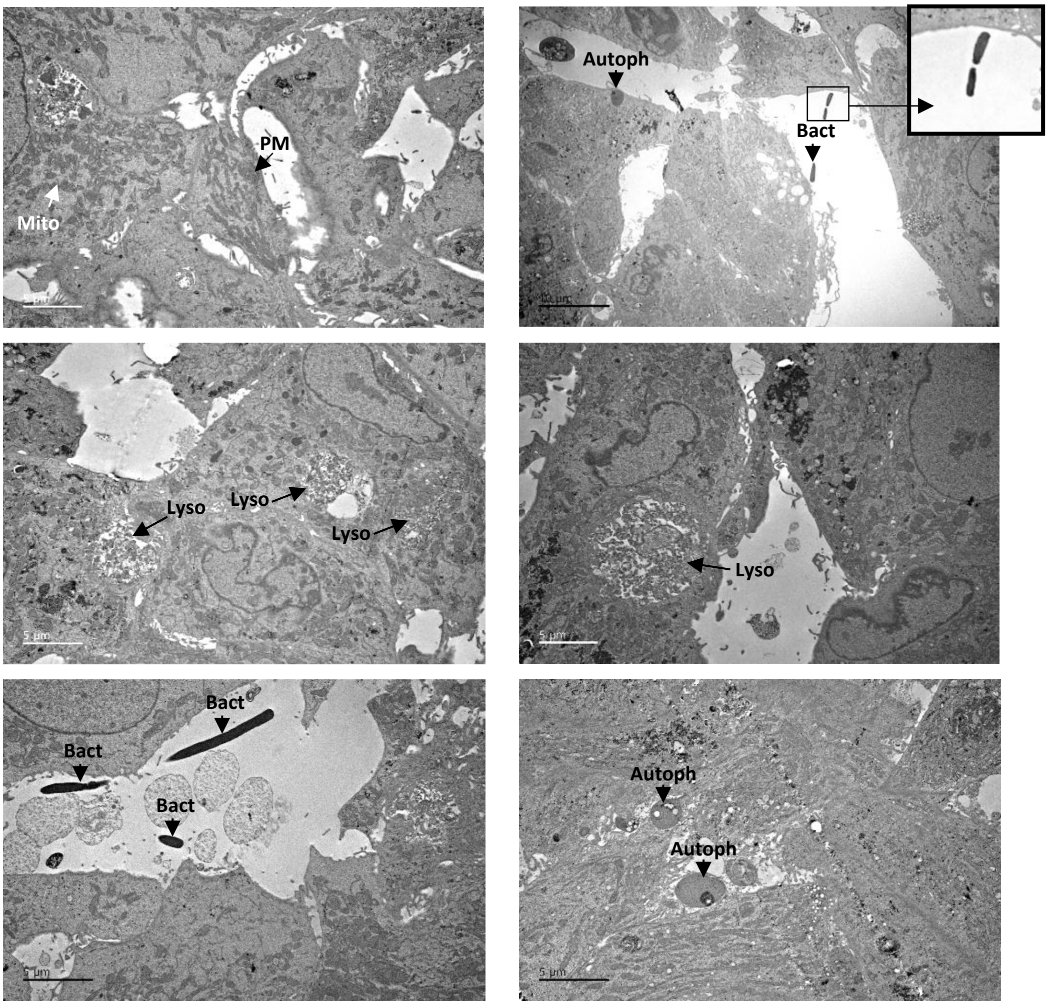


**Supporting Information 8**

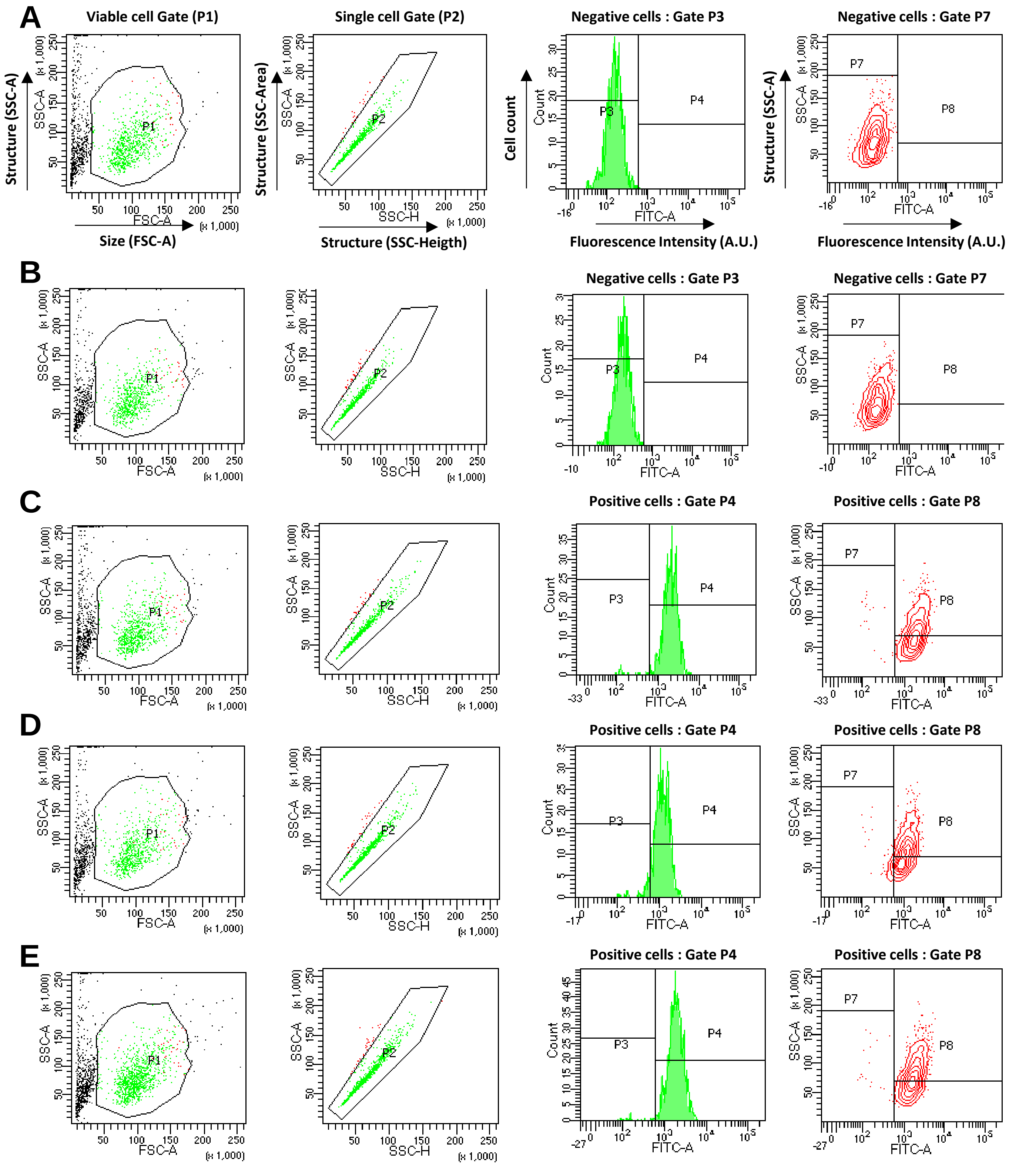


**Supporting Information 9**

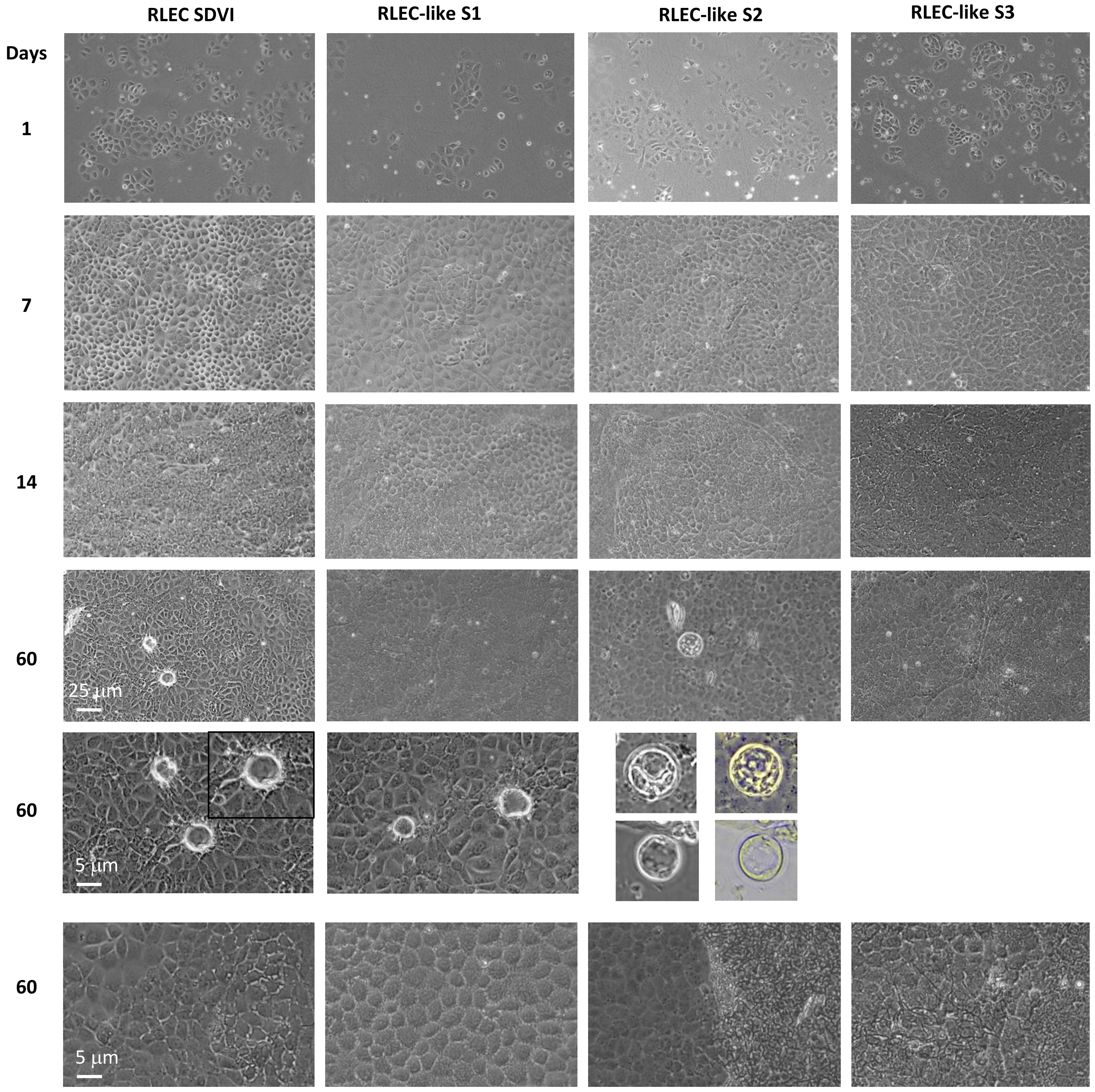


**Supporting Information 10**

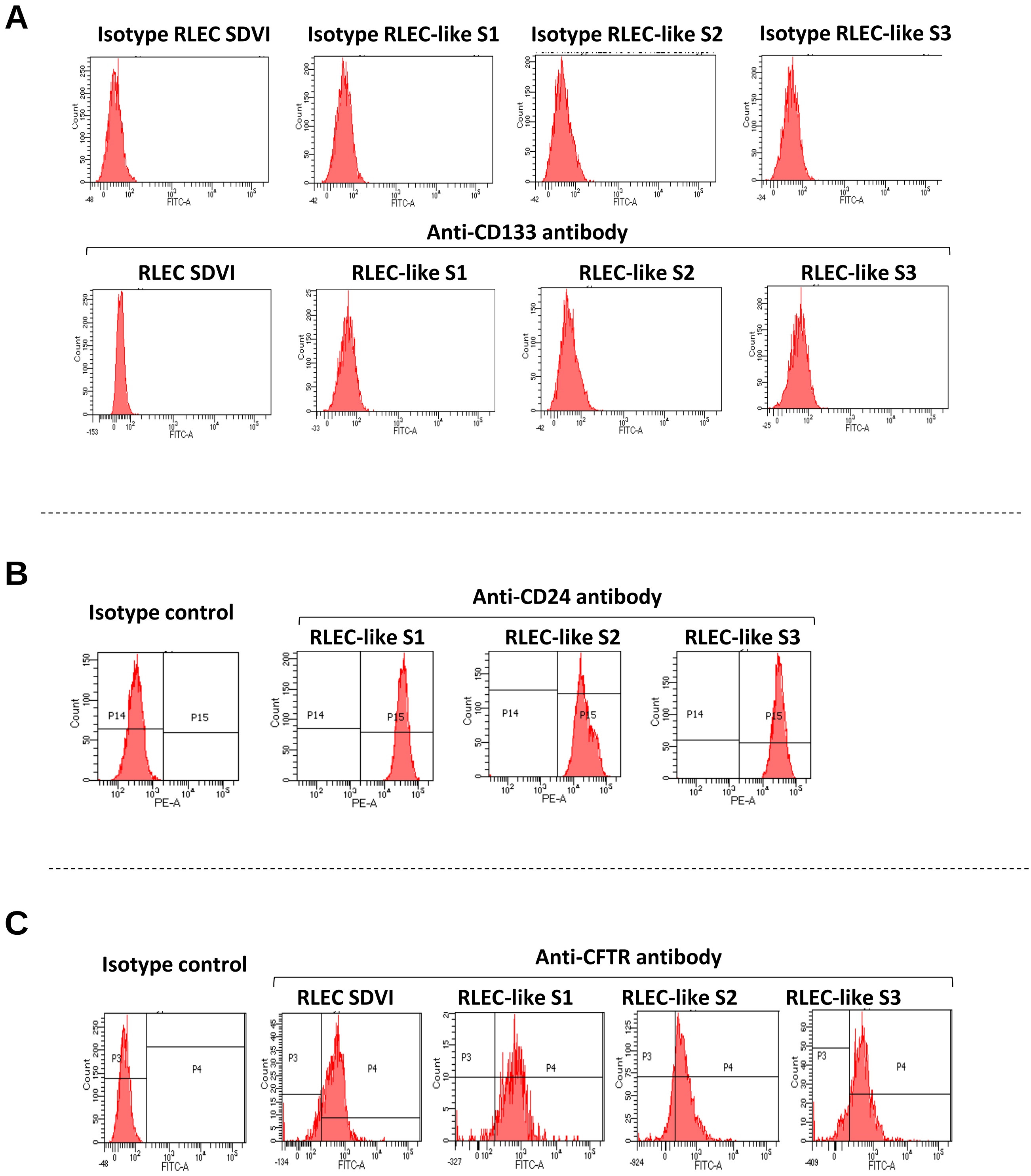


**Supporting Information 11**

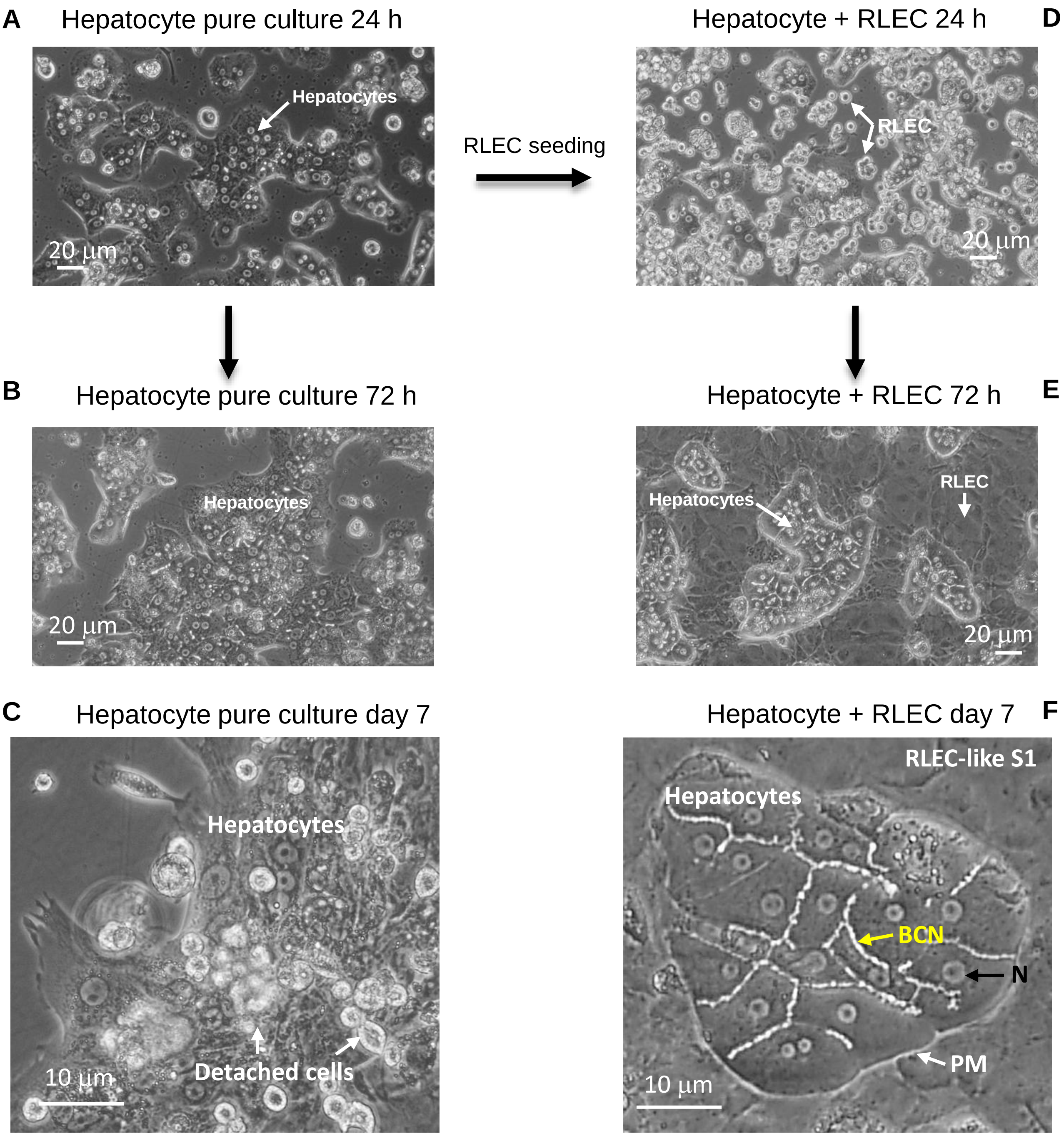


**Supporting Information 12**

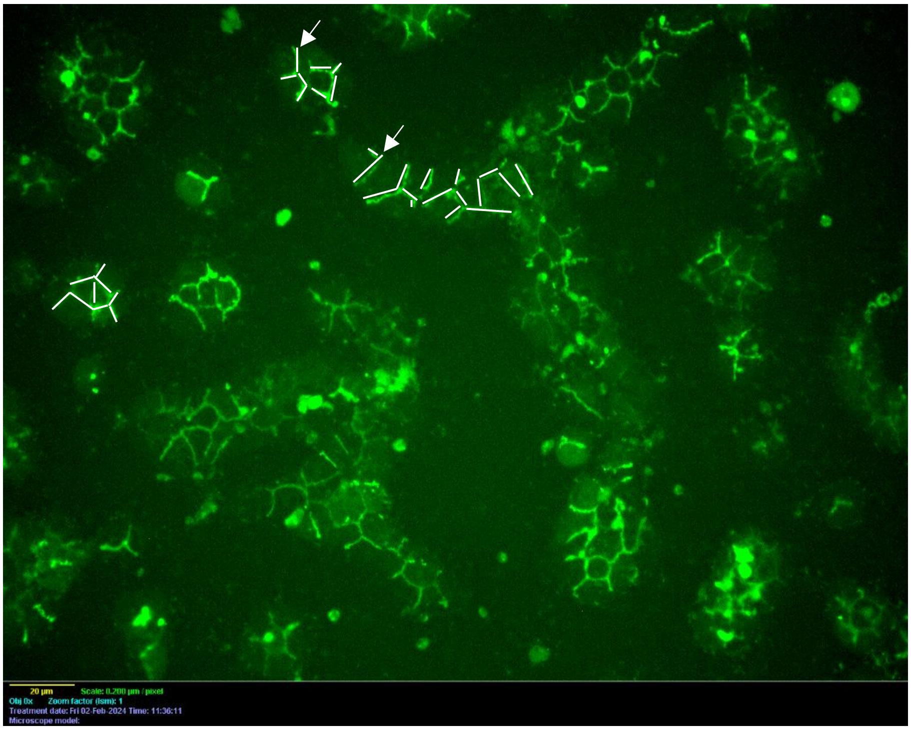


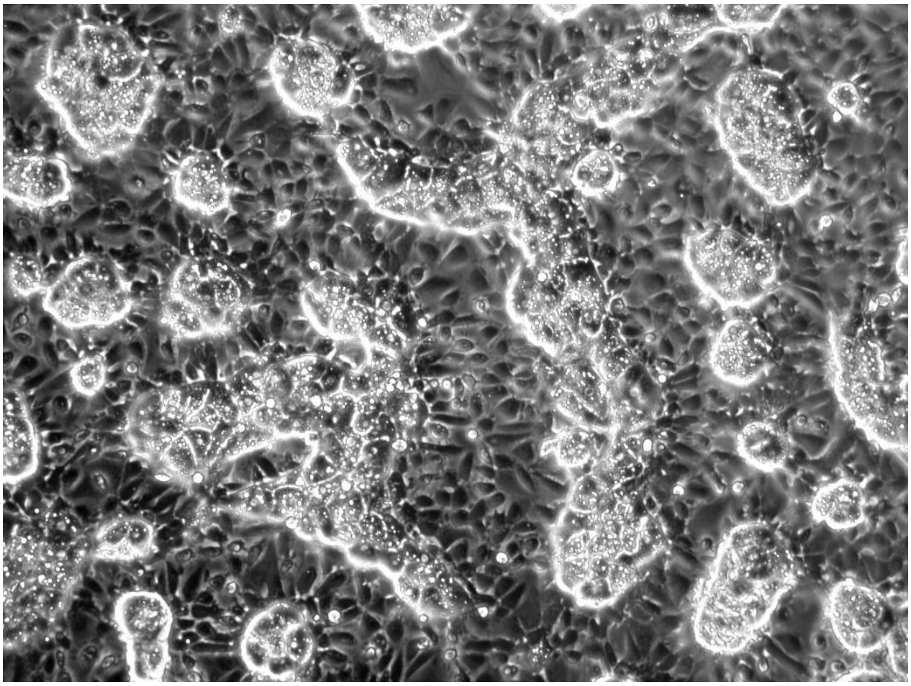

**Supporting Information 13**

| | Forward (5’ to 3’) | Reverse (5’ to 3’) |
|---|---|---|
| Ph.D™-12 sequencing primer | GCCCTCATAGTTAGCGTAACG (single strand sequencing) | |
| 16S rDNA | CTTTCCCTACACGACGCTCTTCCGATCTACTCCTAC GGGAGGCAGCAG (V3F) | GGAGTTCAGACGTGTGCTCTTCCGATCTT ACCAGGGTATCTAATCC (V4R) |
| PsPrSe-16S | GAAGAAGCACCGGCTAACTCCG | ATCTCTACGCATTTCACCGCTAC |
| Albumin | TCAAGAAGGCACCCCGATTA | CTTAGCAAGTCTCAGCAGCAGG |
| Collagen I | GCTTGAAGACCTATGTGGGTATAA | GGGTGGAGAAAGGAACAGAAA |
| TATA Binding Protein | GAGCTGTGATGTGAAGTTTC | TCTGGGTTTGATCATTCTGTA |
| HPRT | GCTTTCCTTGGTCAGGCAGTA | AAGCTT GCGACCTTGACCAT |

**Supporting Information 14**

| Samples | Sex (Male/Female) | Age (years) |
| --- | --- | --- |
| Healthy liver | M | 55 |
| Healthy liver | M | 69 |
| Healthy liver | F | 64 |
| Healthy liver | M | 68 |
| Healthy liver | F | 51 |
| Healthy liver | M | 72 |
| Healthy liver | M | 66 |
| Healthy liver | F | 56 |
| Healthy liver | M | 78 |
| Healthy liver | M | 60 |
| Healthy liver | F | 49 |
| Healthy liver | M | 63 |
| Healthy liver | M | 58 |
| | 9M-4F | 62.2 ± 8.4 |
| Peri-tumoral liver | F | 74 |
| Tumoral liver : CCK | | |
| Peri-tumoral liver | M | 69 |
| Tumoral liver : CCK | | |
| Peri-tumoral liver | M | 74 |
| Tumoral liver : CCK | | |
| Peri-tumoral liver | F | 68 |
| Tumoral liver : CCK | | |
| Peri-tumoral liver | M | 63 |
| Tumoral liver : CCK | | |
| Peri-tumoral liver | M | 55 |
| Tumoral liver : CCK | | |
| Peri-tumoral liver | M | 71 |
| Tumoral liver : CCK | | |
| Peri-tumoral liver | F | 53 |
| Tumoral liver : CCK | | |
| Peri-tumoral liver | M | 57 |
| Tumoral liver : CCK | | |
| Peri-tumoral liver | F | 72 |
| Tumoral liver : CCK | | |
| Peri-tumoral liver | F | 57 |
| Tumoral liver : CCK | | |
| Peri-tumoral liver | F | 60 |
| Tumoral liver : CCK | | |
| Peri-tumoral liver | F | 59 |
| Tumoral liver : CCK | | |
| Peri-tumoral liver | M | 59 |
| Tumoral liver : CCK | | |
| Peri-tumoral liver | M | 61 |
| Tumoral liver : CCK | | |
| Peri-tumoral liver | M | 57 |
| Tumoral liver : CCK | | |
| | 9M-7F | 63 ± 7.1 |

**Supporting Information 15**
